# A *Legionella pneumophila* effector impedes host gene silencing to promote virulence

**DOI:** 10.1101/2022.11.16.516792

**Authors:** Justine Toinon, Monica Rolando, Magali Charvin, Didier Filopon, Lionel Schiavolin, Khadeeja Adam Sy, Hai-Chi Vu, Sarah Gallois-Montbrun, Antoine Alam, Pierre Barraud, Christophe Rusniok, Bérangère Lombard, Damarys Loew, Carmen Buchrieser, Lionel Navarro

**Affiliations:** Institut de Biologie de l’Ecole Normale Supérieure (IBENS), Centre National de la Recherche Scientifique (CNRS), Institut National de la Santé et de la Recherche Médicale (INSERM), Université de recherche Paris, Sciences & Lettres (PSL), 75005 Paris, France; Institut Pasteur, Université de Paris, Biologie des Bactéries Intracellulaires and CNRS UMR 6047, F-75015 Paris, France; Université de Paris, Institut Cochin, INSERM, CNRS, F-75014 Paris, France; Evotec, 40 Avenue Tony Garnier, 69007 Lyon, France; Expression génétique microbienne, Université Paris Cité, CNRS, Institut de biologie physico-chimique, F-75005 Paris, France; Institut Curie, PSL Research University, Centre de Recherche, CurieCoreTech Mass Spectrometry Proteomics, Paris 75248 Cedex 05, France

**Keywords:** *Legionella pneumophila*, type IV effectors, RNA silencing, Argonaute proteins, NF-κB signaling

## Abstract

RNA silencing is a gene silencing mechanism directed by small RNAs. Human miRNAs act as central regulators of host-bacteria interactions. However, it is unknown whether human pathogenic bacteria could impede RNA silencing to promote virulence. Here, we show that the *Legionella pneumophila* type IV-secreted effector LegK1 efficiently suppresses siRNA and miRNA activities in human cells. This effect depends on its known kinase activity, but also on its novel capacity, found here, to bind Argonaute (Ago) proteins. We further demonstrate that the ability of LegK1 to activate NF-κB signaling is required for RNA silencing suppression, establishing a link between effector-mediated NF-κB signaling and RNA silencing suppression. LegK1 also promotes *L. pneumophila* growth in both amoeba and human macrophages, supporting a role for this effector in virulence. Finally, we show that, in infected-macrophages, the latter activity relies, in part, on the genetic targeting of human Ago4. These findings indicate that a *L. pneumophila* effector has evolved to suppress RNA silencing to promote virulence.

**Significance Statement:** It is now well established that mammalian viruses suppress RNAi to promote their replication in host cells. However, whether mammalian pathogenic bacteria use a similar virulence strategy remains unknown. Here, we show that the LegK1 effector from *Legionella pneumophia*, the causal agent of Legionnaires’ disease, efficiently suppresses RNAi in human cells. This effect depends on its ability to interact with Argonaute (Ago) proteins and to activate NF-κB signaling. In addition, LegK1 promotes virulence in infected-macrophages through the genetic targeting of human Ago4. Based on the lack of NF-κB-related factors in amoebae, and on the presence of canonical Ago proteins in these natural *L. pneumophila* hosts, we propose that the RNAi suppression activity of LegK1 represents its primary virulence function.

## Introduction

*Legionella pneumophila* is a Gram-negative bacterium, which infects and replicates in freshwater amoebae and ciliated protozoa in the environment (1, 2). When contaminated water droplets are inhaled by humans, *L. pneumophila* incidentally infects lung alveolar macrophages, which can result in a severe form of pneumonia called Legionnaires’ disease (2, 3)*. L. pneumophila* encodes a type IV secretion system, referred to as the Dot/Icm system, to promote intracellular replication (4, 5). This secretion system translocates over 330 effectors into host cells and is essential for the replication of *L. pneumophila* in protozoan hosts and in human macrophages (6, 7). A significant proportion of *L. pneumophila* effectors resembles eukaryotic-like proteins or encode domains preferentially present in eukaryotic proteins, which have likely been co-opted during co-evolution with their protozoan hosts to help subverting host functions (7–10). Among them, is the family of LegK proteins that encodes eukaryotic-like kinase domains targeting specific host proteins (10). For example, LegK1 is a serine/threonine protein kinase that phosphorylates the NF-κB inhibitor IκBα, as well as other IκB and NF-κB family members, resulting in a potent NF-κB signaling activation in human cells (11). This effector contributes to the sustained Dot/Icm-dependent NF-κB signaling detected during *L. pneumophila* infection of human alveolar macrophages (12–14). Furthermore, it triggers the induction of anti-apoptotic and host cell survival genes (12–14), which is thought to promote *L. pneumophila* growth in human macrophages.

MicroRNAs (miRNAs) are small non-coding RNAs (sRNAs) that post-transcriptionally regulate genes through the targeting of sequence complementary mRNA targets. In mammals, miRNA biogenesis involves the cleavage of primary miRNA transcripts by the Drosha-DGCR8 complex, which further releases hairpin-based miRNA precursors that are subsequently processed by the RNAse III enzyme Dicer (15). The resulting miRNA duplexes then bind to an Argonaute (Ago) protein, and one strand, the guide, remains associated to this silencing factor to form a miRNA-Induced Silencing Complex (miRISC) (16). The miRISC further directs silencing of mRNA targets through endonucleolytic cleavage (slicer activity), translational repression and/or mRNA degradation (16). Four structurally conserved Ago proteins (Ago1-4) are encoded by the human genome (17). Human Ago2 is the most characterized Ago member that possesses a slicer activity directed by a catalytic tetrad, DEDH (Asp-Glu-Asp-His), embedded in its PIWI domain (17). This factor is the central component of the miRISC. Mechanistically, the phosphorylation of Ago2 at serine 387 triggers its binding to trinucleotide repeat containing 6 (TNRC6) proteins, which are tryptophan (W)-rich cofactors that associate with the PIWI domain of Ago2 through W-repeats (W-motifs) found in their N-terminal Ago-binding domains (ABDs) (18, 19). These molecular events are essential for High Molecular Weight (HMW)-miRISC assembly by recruiting downstream effector proteins, including those involved in decapping, and deadenylation processes (20). In addition, TNRC6 proteins use W-motifs embedded in their C-terminal silencing domains to directly bind CCR4-NOT and PAN2-PAN3 (poly(A)-specific ribonuclease 2-3) deadenylase complexes, thereby contributing to HMW-miRISC activity (20). This series of molecular events leads to the translational repression, destabilization and degradation of miRNA targets, and in turn regulates diverse biological processes, including development (*e.g.* neurological development), differentiation, stress signaling and antiviral responses (21–25). Human Ago1 and Ago4 are slicer-deficient and thus trigger gene silencing through slicer-independent mechanisms (26). In contrast, human Ago3 retains a canonical DEDH catalytic tetrad and was recently shown to direct RNA cleavage when loaded with 14-nt long guide miRNAs (17, 27). Although little is known about the biological functions of these three Ago proteins, gene deletions of *Ago1* and *Ago3* have been retrieved in patients suffering from neurological disorders (17, 28). Furthermore, several biological functions have been ascribed to mammalian Ago4. This factor (i) regulates meiotic entry and meiotic sex chromosome inactivation in mice germ cells (29), (ii) translationally inhibits the second cistron of the *CACNA1A* locus in a miRNA-dependent manner, thereby inhibiting the growth of neurons and Purkinje cells in mice (30), (iii) interacts with the *de novo* methyltransferase DNMT3A and in turn directs cytosine methylation of some mature miRNAs, which inhibits their functions and is associated with poor prognosis in glioma (31), (iv) mediates RNA-dependent DNA methylation in human cells (32), and (v) promotes antiviral resistance in mice (33). Therefore, despite early assumptions that mammalian Ago proteins act redundantly, they can also exhibit very specialized functions.

Human miRNAs have been extensively studied in the context of host-bacteria interactions (34). They regulate various cellular processes during bacterial infection, including innate immune responses, host cell cycle, cell death and survival pathways, autophagy, and cytoskeleton organization (34). For example, *L. pneumophila* triggers the differential expression of 85 human miRNAs during infection of macrophages (35). Among them, a trio of down-regulated miRNAs, namely miR-125b, miR-221 and miR-579, was shown to target repressors of *L. pneumophila* replication in macrophages (35). Interestingly, *L. pneumophila* infection is not only modulating the expression of human miRNAs, but *L. pneumophila* is itself translocating active small RNAs (sRNAs) *via* extracellular vesicles (EVs) into the host cell (36). Indeed, 15 sRNAs were identified that mimic human miRNAs and two of these, namely RsmY and tRNA-Phe, down-regulate selected sensor and regulator proteins of the host innate immune response in a miRNA-like manner (36). Intriguingly, a recent study also shows that human airway cells make use of EVs to deliver let-7b-5p into *Pseudomonas aeruginosa* and reprogram bacterial gene expression, thereby reducing biofilm formation and antibiotic resistance (37). Altogether, these studies, among many others, highlight a central role played by sRNAs in the reprogramming of both host and bacterial gene expression during infections.

Although the role of human miRNAs in host-bacteria interactions is now well-established, there is currently no evidence indicating that human pathogenic bacteria can interfere with RNA silencing as part of their virulence functions. This has, however, been previously characterized in a phytopathogenic *Pseudomonas syringae* strain, which delivers several type III-secreted effectors into host cells to suppress different steps of the Arabidopsis miRNA pathway (38). It has also been extensively characterized in mammalian viruses that produce Viral Suppressors of RNA silencing (VSRs) to replicate in host cells (39–41).

Here, we show that the *L. pneumophila* type IV-secreted effector LegK1 acts as a *bona fide* Bacterial Suppressor of RNA silencing (BSR). We demonstrate that LegK1 efficiently suppresses siRNA- and miRNA-activities in human cells through both its kinase activity and a newly-established Ago-binding capacity. We further demonstrate that the ability of LegK1 to activate NF-κB signaling is required for RNA silencing suppression. In addition, we found that LegK1 promotes *L. pneumophila* growth in both human macrophages and amoeba, supporting a virulence function of this effector in biologically relevant host cells. Importantly, the genetic targeting of human Ago4 by LegK1 was found to promote the growth of *L. pneumophila* in infected-macrophages. Therefore, besides identifying the first BSR from a human pathogenic bacterium, this study unveils an unexpected link between effector-mediated NF-κB signaling activation and RNA silencing suppression. It also provides evidence that a *L*. *pneumophila* effector has evolved to suppress RNA silencing as part of its virulence function.

## Results

### The LegK1 effector suppresses siRNA-directed silencing in human cells

The known Ago-binding function of W-motifs, present in some endogenous silencing factors as well as in some VSRs (20, 39, 42), prompted us to examine whether effectors from human pathogenic bacteria have also evolved such Ago-binding platforms to interfere with RNA silencing. To test this hypothesis, we retrieved the protein sequences from 25 genomes of pathogenic bacteria and used them with the Wsearch algorithm to retrieve putative W-motifs (43) (*SI Appendix*, Fig. S1*A*). The score of each W-motif was determined using the animal Bidirectional Position-Specific Scoring Matrix (BPSSM), generated from experimentally-verified Ago-binding proteins and their orthologous sequences (43). Among the candidate proteins exhibiting the highest W-score, we selected a subset of secreted virulence factors and tested their possible effects on the siRNA-based *CXCR4* reporter (44) (*SI Appendix*, Fig. S1*B*). This reporter system relies on the co-transfection of one plasmid expressing the *firefly luciferase* fused to the 3’UTR of *CXCR4,* a second one expressing the *renilla luciferase*, which served as a transfection control, along with exogenous siRNAs targeting the 3’UTR of *CXCR4*. We first showed that the silencing of this *firefly luciferase* reporter was compromised in the *ago2*^-/-^ and *ago1*^-/-^/*ago2*^-/-^ CRISPR/Cas9-based mutant HeLa cell lines, as revealed by an increased luminescence intensity compared to control cells (*SI Appendix*, Fig. S1*C* and *D*). In contrast, the silencing of the *firefly luciferase* remained effective in *ago1*^-/-^ and *dicer*^-/-^ cell lines. Collectively, these results indicate that the *CXCR4* reporter system is dependent on Ago2, but not on Ago1. They also indicate that Dicer is not required for this process, which is consistent with former studies showing that sRNA duplexes can be loaded in mammalian Ago2 in a Dicer-independent manner (16). Significantly, when monitoring luminescence intensity in HeLa cells expressing individually these 8 candidate bacterial W-effectors (*SI Appendix*, Fig. S1*E*-*G*), we found that LegK1 from *L. pneumophila* (strain Paris) was the sole effector able to inhibit the silencing of the *CXCR4* reporter, as found in *ago2*^-/-^ cells or in cells expressing the VSR *Hepatitis B virus* HBx (45) (*SI Appendix*, Fig. S1*H* and *I*). Altogether, these data indicate that LegK1 suppresses the silencing of an Ago2-dependent siRNA-based reporter system.

### LegK1 suppresses RNA silencing through its kinase activity and W-motifs

LegK1 is an experimentally validated Dot/Icm type IV-secreted effector possessing a eukaryotic-like serine/threonine kinase activity (11). Here, we additionally found that LegK1 contains a putative Ago-binding platform composed of four predicted W-motifs (*SI Appendix*, Fig. S1*J*). The tryptophan residues located at positions 41, 283 and 293 of the LegK1 protein sequence, namely W41, W283 and W293, displayed the highest W-score, and were thus selected for further analyses (Fig. 1*A* and *SI Appendix*, Fig. S1*J*). We first substituted each of these three tryptophans by phenylalanine (W>F; LegK1-3WF). An *in*-*silico* analysis indicated that these point mutations have a very limited destabilization effect (*SI Appendix*, Fig. S2*A*). In parallel, we generated a K121A substitution in the LegK1 ATP-binding site (LegK1-KA), which has been previously shown to abolish the kinase activity of this effector (11). Both LegK1-3WF and LegK1-KA proteins accumulated to the same levels as wild-type (WT) LegK1 proteins in HEK293T cells (Fig. 1*B*), indicating that these mutated proteins are stable when expressed in human cells. We further analyzed their ability to suppress RNA silencing. To this end, we used a siRNA-based *GFP* reporter system that exhibits less variability in human cells, as compared to the *CXCR4* reporter, because it only relies on the co-transfection of a plasmid expressing the *enhanced GFP* (*eGFP*), together with anti-*GFP* siRNAs targeting a single site located in the 5’ region of the *eGFP* coding sequence. Furthermore, this siRNA-based reporter system has been previously used to characterize VSRs from different human pathogenic viruses (46). We found that this reporter was robustly silenced by the anti-*GFP* siRNAs, as revealed by low accumulation of eGFP proteins when expressed in HeLa and HEK293T cells (Fig. 1*C*-*E*). In contrast, the eGFP protein accumulation was moderately increased in the *ago2*^-/-^ HeLa cell line expressing the anti-*GFP* siRNA duplexes, an effect that was found enhanced in the *ago1*^-/-^/*ago2*^-/-^ mutant cells (Fig. 1*C*). These results indicate that both Ago1 and Ago2 proteins direct the silencing of this *GFP*-based reporter system. Of note, the fact that the reporter was not fully derepressed in *ago1*^-/-^/*ago2*^-/-^ HeLa cells suggests that Ago3 and/or Ago4 must additionally contribute to its silencing. When the same analysis was conducted in HEK293T cells expressing LegK1, we found an efficient suppression of *eGFP* silencing, a process that was not observed in cells expressing the empty vector or the type III-secreted YopM effector from *Yersinia pestis*, which is deprived of silencing suppression activity (Fig. 1*D* and *E*). It is noteworthy that the silencing suppression effect mediated by LegK1 was even more pronounced than the one triggered by T6B, a TNRC6B-derived peptide containing multiple W-motifs, which was shown to disrupt the function of multiple Ago proteins (47) (Fig. 1*E*). Taken together, these data indicate that LegK1 efficiently suppresses the silencing of another siRNA-based reporter controlled by multiple Ago proteins. Importantly, we found that the LegK1-3WF and LegK1-KA mutant versions no longer triggered silencing suppression of this siRNA-based *GFP* reporter (Fig. 1*E*). Therefore, both the kinase activity and the putative Ago-binding platform of LegK1 are essential for the silencing suppression of this siRNA-based reporter.

**Fig. 1.**
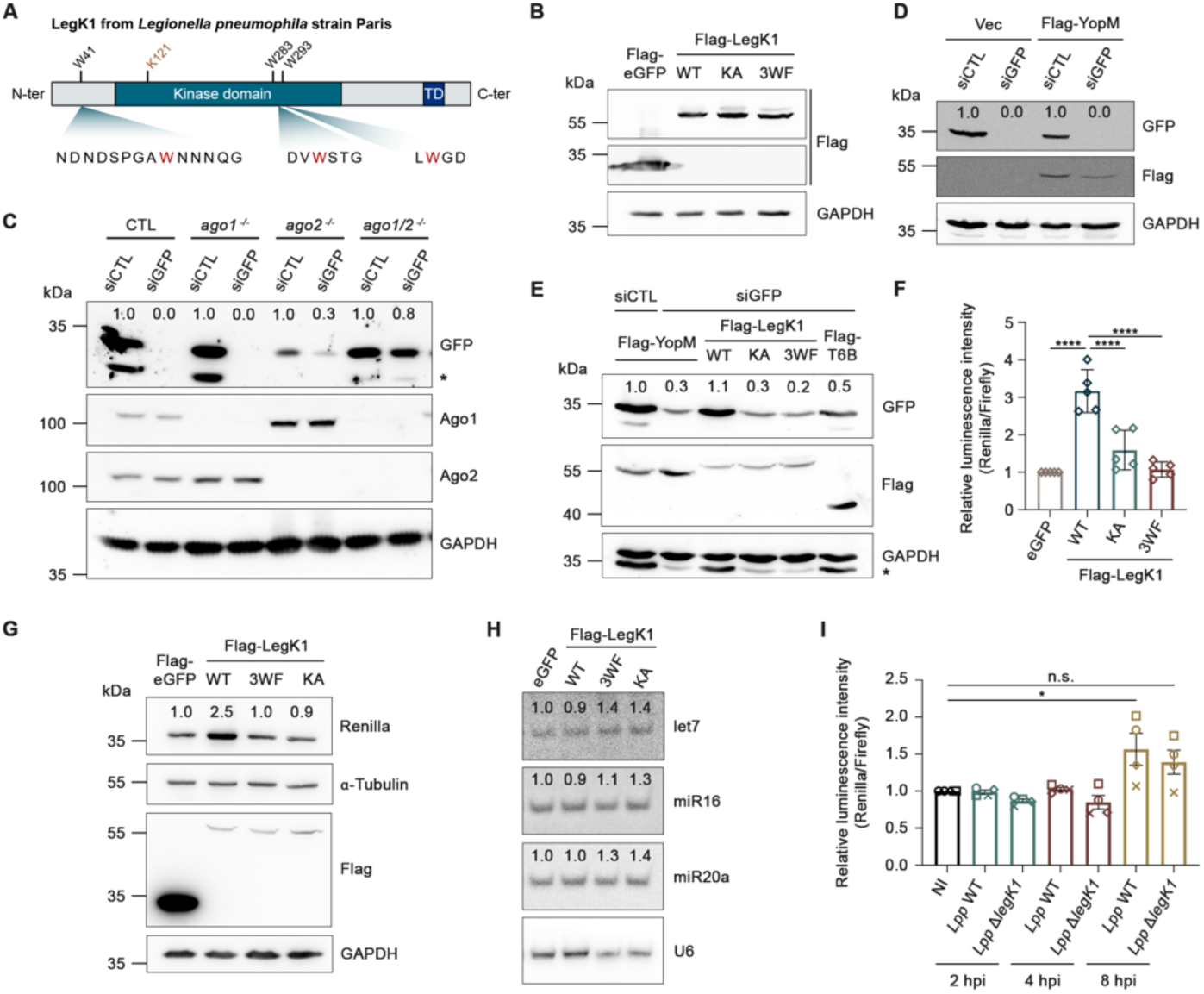
LegK1 suppresses siRNA- and miRNA-guided gene silencing through both its kinase activity and its predicted Ago-binding platform. (*A*) Schematic representation of the LegK1 protein sequence. The position and amino acid sequence of the three selected W-motifs (W41, W283, W293) recovered from the Wsearch analysis are depicted. The lysine 121 is located inside the ATP-binding pocket of the serine/threonine kinase of LegK1 and is required for catalytic activity. TD, transmembrane domain. (*B*) The LegK1-KA and LegK1-3WF mutant proteins are stable in human cells. HEK293T cells were transfected with vectors expressing either Flag-eGFP, 2xFlag-LegK1 (WT), 2xFlag-LegK1-KA or 2xFlag-LegK1-3WF proteins for 48 hours. Immunoblot was performed using an anti-Flag antibody. (*C*) A GFP-based siRNA reporter is silenced by both Ago1 and Ago2. Control (CTL), *ago1^-/^*^-^, *ago2^-/^*^-^, *ago1/2^-/^*^-^, and *dicer^-/^*^-^ HeLa cell lines were co-transfected with vectors expressing eGFP and control siRNAs (siCTL) or GFP siRNA duplexes (siGFP). At 48h post-transfection, cell lysates were subjected to Western Blot analysis with indicated antibodies. The eGFP protein levels are relative to the ones of GAPDH and further normalized to the corresponding siCTL condition, as indicated above the GFP blot. * represents a specific band. (*D*) YopM does not alter the silencing of the GFP-based siRNA reporter. GFP-based silencing reporter assay in the presence of empty vector (Vec) or YopM in HEK293T cells. The experiment and the eGFP protein quantification were performed as described in (*C*). (*E*) LegK1 suppresses the silencing of a GFP-based siRNA reporter through both its kinase activity and W-motifs. GFP-based silencing reporter assay in the presence of LegK1 WT, LegK1-KA or LegK1-3WF in HEK293T cells. The experiment was performed as described in (*C*), but the vector expressing Flag-YopM, 2xFlag-LegK1-KA, 2xFlag-LegK1-3WF or Flag-T6B were co-transfected along with the reporter system. * represents aspecific bands. (*F*) LegK1 suppresses the silencing of a let-7a miRNA luciferase reporter through both its kinase activity and W-motifs. HEK293T cells were co-transfected with let-7a luciferase reporter and vectors expressing Flag-eGFP, 2xFlag-LegK1, 2xFlag-LegK1-KA or 2xFlag-LegK1-3WF. The luciferase expression was measured and normalized to the eGFP condition. (*G*) The cell lysates from the let-7a miRNA luciferase reporter assay were subjected to Western Blot analysis with indicated antibodies. The Renilla protein levels are relative to the ones of GAPDH and further normalized to the corresponding eGFP condition, as indicated above the Renilla blot. (*H*) LegK1 does not alter the accumulation of mature let-7a miRNA. Total RNAs of HEK293T cells expressing eGFP, LegK1 WT, LegK1-3WF or LegK1-KA were extracted and subjected to a Northern blot analysis with indicated probes. The miRNA levels are relative to the ones of U6 miRNA and further normalized to the eGFP condition. (*I*) The *Lpp* strain derepresses let-7a luciferase reporter in HEK293-FcɣRII cells, partially through LegK1 effector. Cells were co-transfected with let-7a luciferase reporter for 24h and were infected with *Lpp* WT strain at an MOI of 10. Luciferase expression was measured at 2, 4 and 8h post-infection. The luminescence intensity was measured as in (*F*). NI corresponds to non-infected. All data are representative of at least three independent experiments. GAPDH and α-tubulin were used as loading controls in all these experiments. *, *P*< 0.05; ****, P< 0.0001; n.s., not significant, as calculated using ordinary one-way ANOVA (Dunnett’s multiple comparisons test).

To investigate whether LegK1 additionally suppresses miRNA functions, we made use of a miRNA-based luciferase reporter system containing bulged target sites. More specifically, we generated a plasmid composed of two constructs: a non-targeted *firefly luciferase*, which served as an internal reference control, and a miRNA-targeted *renilla luciferase* carrying, downstream of its coding sequence, a let-7a complementary sequence containing mismatches opposite to nucleotides at positions 9, 10 and 11 of the let-7a sequence (44). We found that the co-expression of LegK1 with the *let-7a* reporter led to an enhanced Renilla luciferase protein accumulation and luminescence intensity as compared to the co-expression of eGFP with this reporter system (Fig. 1*F* and *G*). In contrast, these effects were not detected in HEK293T cells expressing LegK1-KA and LegK1-3WF mutants. Collectively, these data demonstrate that LegK1 additionally inhibits the function of a well-characterized endogenous miRNA through both its kinase activity and a predicted W-dependent Ago-binding platform. This LegK1-triggered molecular effect is not caused by a decrease in the accumulation of the mature miRNA let-7a, as revealed by Northern blot analysis (Fig. 1*H*). Furthermore, LegK1 does not interfere with the accumulation of two other canonical human miRNAs, namely miR16 and miR20a (Fig. 1*H*). Therefore, LegK1 likely acts at the level of miRNA activity rather than on miRNA biogenesis or stability. Altogether, these data indicate that LegK1 suppresses siRNA and miRNA activities through both its kinase activity and a predicted Ago-binding platform.

### LegK1 dampens let-7a function in a physiological context of infection

Having shown that LegK1 acts as a silencing suppressor upon artificial transfection of this effector in HEK293T cells, we next examined whether this novel activity could be detected during infection. To test this hypothesis, we generated an isogenic *legK1* deletion mutant, referred to here as the *Lpp* Δ*legK1* strain. Whole-genome sequencing of *Lpp* Δ*legK1* confirmed a unique deletion in the *legK1* gene. This deletion did not affect the fitness of this bacterium *in vitro* (*SI Appendix*, Fig. S3). We further transfected HEK293 cells stably expressing the FcɣRII receptor (48), with the above *let-7a* sensor, and further infected these cells with either the *Lpp* WT or Δ*legK1* strains. Significantly, and despite the unlikely infection of all the cells expressing the *let*-*7a* reporter system, we found that the *Lpp* WT strain derepressed this miRNA-dependent sensor at 8 hours post-infection (hpi) (Fig. 1*F* and *I*). These data provide thus evidence that *L*. *pneumophila* suppresses let-7a function in a physiological context of infection. Furthermore, the derepression of the let-7a reporter system appeared lower in response to *Lpp* Δ*legK1*, compared to the WT strain at this infection timepoint (Fig. 1*I*). Altogether, these data demonstrate that *L*. *pneumophila* dampens let-7a function during infection and that this effect might be in part achieved by LegK1.

### LegK1 interacts with Ago2, Ago4 and other miRISC factors in human cells

The presence of predicted W-motifs in the protein sequence of LegK1 and their relevance in silencing suppression, suggested that this effector could interact with human Ago proteins. To test this possibility, we expressed the Flag-tagged LegK1-WT, LegK1-KA and LegK1-3WF versions in HEK293T cells and immunoprecipitated their cellular partners. Using Western blot analysis, we demonstrated that Ago1, Ago2 and Ago4 proteins were recovered from Flag-LegK1 immunoprecipitates, which was not observed after immunoprecipitation of Flag-eGFP from control cells (Fig. 2*A* and *B*). This result indicates that LegK1 interacts with protein complexes containing Ago1, Ago2 and Ago4 proteins. It is also consistent with the co-localization found between a mCherry-LegK1 fusion and GFP-Ago2 or GFP-Ago4 in HEK293T cells, which was not detected between mCherry and GFP-Ago2 or GFP-Ago4 (Fig. 2*C*-*E*), nor between mCherry-LegK1 and GFP (*SI Appendix*, Fig. S4), which served as negative controls. Interestingly, we also recovered in the above Flag-LegK1 immunoprecipitates the TNRC6A, PABPC1 and DDX6 proteins (Fig. 2*A* and *B*), which are well-characterized HMW-miRISC factors (20). This suggests that LegK1 interacts with mature Ago-RISCs engaged in RNA target repression. Importantly, the above interactions were maintained with the catalytic mutant LegK1-KA, indicating that the kinase activity of LegK1 is dispensable for this process. In contrast, they were significantly reduced with the LegK1-3WF mutant (Fig. 2*A* and *B*), despite unaltered effects on the co-localization of the mCherry-LegK1-3WF fusion protein with GFP-Ago2 and GFP-Ago4 (Fig. 2*C*). Collectively, these data indicate that the Ago-binding platform of LegK1 is required to interact with Ago-RISCs, but it does not interfere with the capacity of LegK1 to co-localize with Ago2 and Ago4 in human cells. Finally, we found that RNase A treatment did not alter the ability of Flag-LegK1 to interact with GFP-Ago2 (Fig. 2*F*), demonstrating that LegK1 interacts with Ago2 in an RNA-independent manner.

**Fig. 2.**
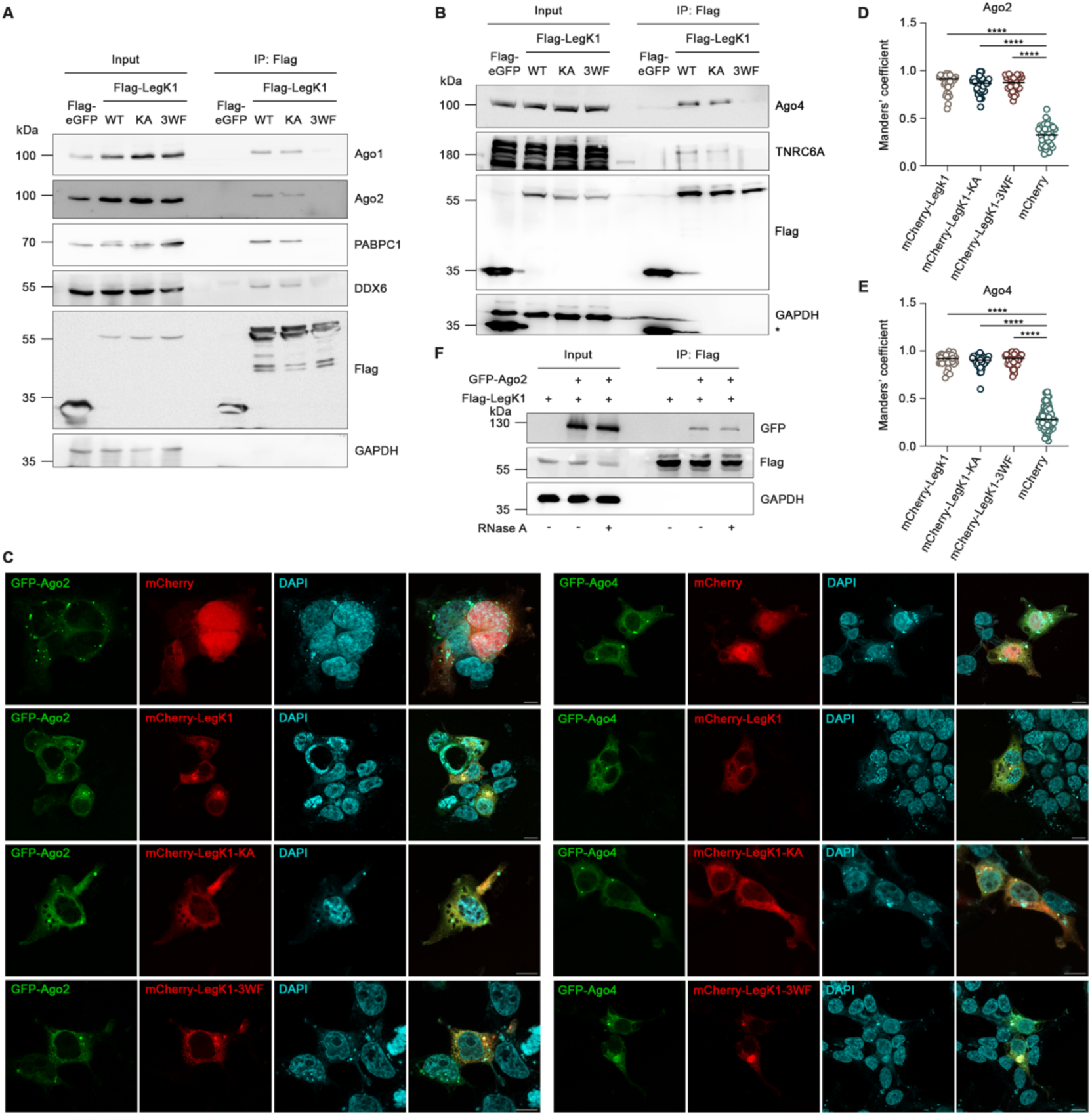
LegK1 interacts with Ago1, Ago2 and Ago4-containing miRISCs. (*A*) and (*B*) LegK1 interacts with Ago proteins and components of the miRISC through its W-motifs. HEK293T cells were transfected with vectors expressing Flag-eGFP, 2xFlag-LegK1 WT, 2xFlag-LegK1-KA or 2xFlag-LegK1-3WF. At 48h post-transfection, cells were lysed and proteins were immunoprecipitated using an anti-Flag antibody. Total cell lysates (input) and the immunoprecipitates (IP: Flag) were analyzed by immunoblotting using indicated antibodies. * represents a specific band. (*C*) LegK1 co-localizes with Ago2 and Ago4 in HEK293-FcɣRII cells. Representative confocal microscopy images of HEK293-FcγRII cells transfected for 48 hours with eGFP-Ago2 and eGFP-Ago4 together with mCherry-LegK1 and its mutated forms, as indicated. Empty peGFP-C1 and pmCherry-C1 were used as negative controls. DAPI and Phalloidin are in cyan. Single channel and merged images are shown. Z stack images were acquired and deconvolution images are presented as maximum intensity z-projections. Scale bars 10 µm. (*D*) and (*E*) Quantification of colocalizations in (*C*). The fraction of mCherry-LegK1 and its mutated forms that overlaps with GFP-Ago2 (*D*) or GFP-Ago4 (*E*) was quantified by Manders’ colocalization coefficient from 3 independent experiments. (*F*) LegK1 interacts with Ago2 in an RNA-independent manner. HEK293T cells were transfected with a vector expressing 2xFlag-LegK1 or co-transfected with vectors expressing 2xFlag-LegK1 and GFP-Ago2. At 48h post-transfection, cells were lysed, incubated with or without RNase A, and proteins were further immunoprecipitated using an anti-Flag antibody. Total cell lysates (input) and the immunoprecipitates (IP: Flag) were analyzed by immunoblotting using indicated antibodies.

### LegK1 interacts with human Ago2 and its PIWI domain *in vitro*

We next investigated whether LegK1 directly interacts with human Ago2. We conducted an *in vitro* interaction assay using GST-Ago2 and His6-SUMO-LegK1^2:386^ –deprived of its predicted transmembrane domain–, which were both expressed and purified from *E. coli* (Fig. 3*A* and *B*, *SI Appendix*, Fig. S5). We found that the GST-tagged Ago2 protein efficiently bound to His6-SUMO-LegK1^2:386^, while it did not interact with the TAP negative control (Fig. 3*B*). Because the PIWI domain of human Ago2 is known to possess W-binding interfaces (20), we additionally expressed and purified a CBP-tagged PIWI domain of human Ago2 (Fig. 3*A* and *C*). Using an *in vitro* pull-down assay, we found that the CBP-PIWI^517:859^-His6 recombinant protein interacts with His6-SUMO-LegK1^2:386^, however this interaction was not higher in the presence of RNA extracts from HEK293T cells (Fig. 3*C*). Therefore, LegK1 directly interacts with human Ago2, likely through interaction surfaces embedded in its PIWI domain. Our data also suggest that one or several W-motifs of LegK1 might orchestrate the interaction between this bacterial effector and human Ago proteins.

**Fig. 3.**
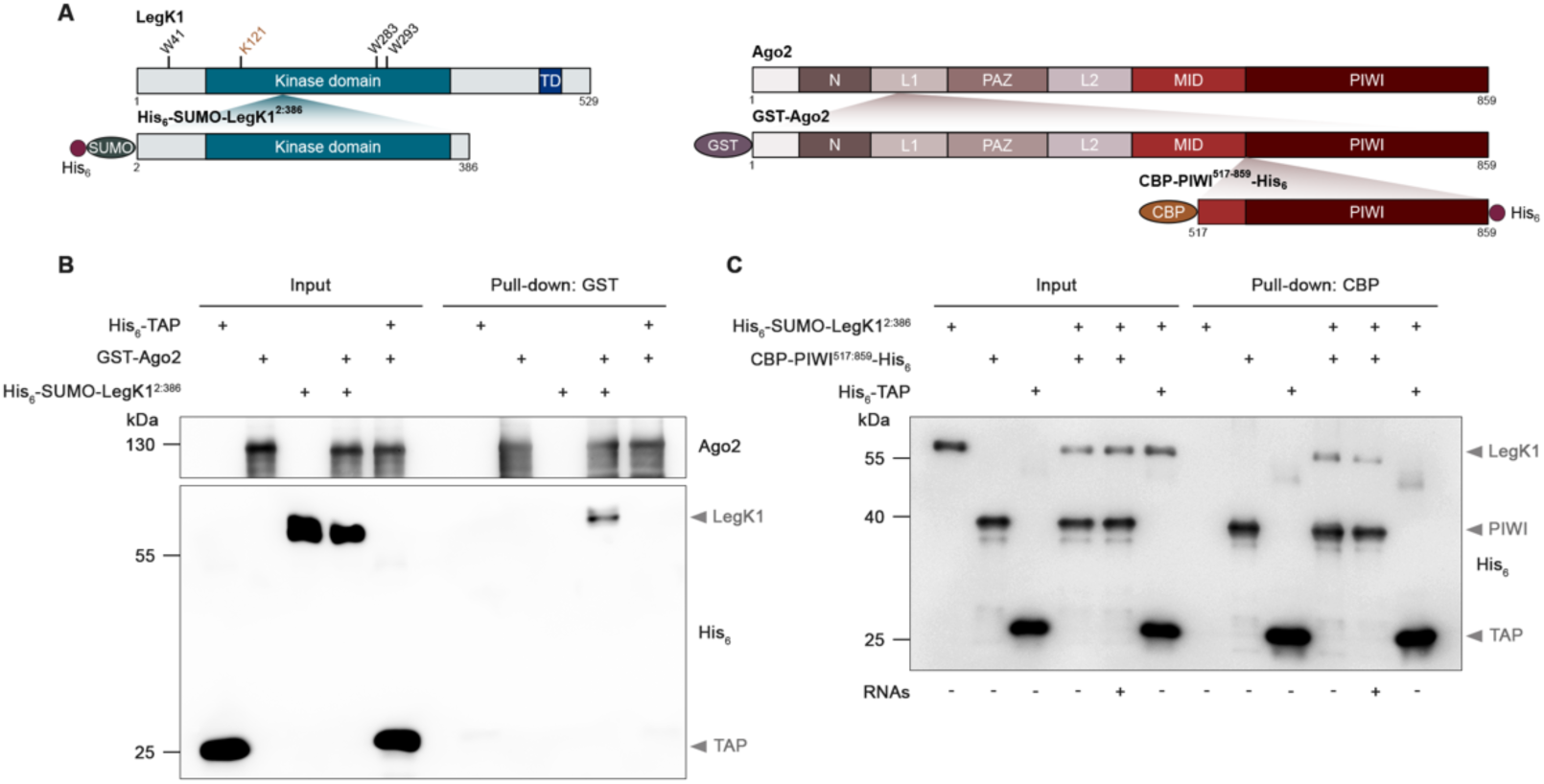
LegK1 directly interacts with Ago2 and its PIWI domain. (*A*) Schematic representation of the recombinant proteins used for *in vitro* pull-down assays. CBP, Calmodulin Binding Protein; GST, Glutathione S-Transferase; SUMO, Small Ubiquitin-like MOdifier; TD, Transmembrane Domain. (*B*) LegK1 directly interacts with Ago2 *in vitro*. Pull-down assay between human Ago2 and LegK1 recombinant proteins. GST-Ago2 was incubated with His6-SUMO-LegK1 (a.a. 2:386) or His6-TAP used as negative control, and subjected to a GST pull-down assay. (*C*) LegK1 directly interacts with the PIWI domain of Ago2 *in vitro.* Pull-down assay between the PIWI domain of human Ago2 and LegK1 recombinant proteins in the absence or presence of RNAs. CBP-PIWI-His6 (a.a. 517-817) or His6-TAP, which consists of a CBP tag used as negative control, were incubated with His6-SUMO-LegK1 (a.a. 2:386), with or without RNAs extracted from human cells, and subjected to a CBP pull-down assay. All data are representative of three independent experiments. GAPDH was used as a loading control.

### The W-motifs of LegK1 exhibit Ago-binding capacity *in vitro*

To further examine the contribution of the selected W-motifs of LegK1 in Ago-binding, we first attempted to express and purify the LegK1-3WF mutant version from *E. coli*. However, this mutant formed insoluble inclusion bodies, preventing further *in vitro* pull-down experiments (*SI Appendix*, Fig. S5). To circumvent this issue, we chemically synthesized biotinylated peptides containing each candidate W-motif (W41, W283 and W293), surrounded by native amino acid residues, referred to here as W1, W2 and W3 peptides (Fig. 4*A*). In parallel, we synthetized identical peptides where each tryptophan was substituted by phenylalanine, named F1, F2 and F3 mutant peptides. A pull-down experiment of each individual peptide was performed to elute bound proteins. Using this approach, we found that the W1 peptide did not bind to any of the Ago proteins, indicating that W41 is not a functional Ago-binding motif (Fig. 4*B*). In contrast, the W2 and W3 peptides readily bound to human Ago1, Ago2, Ago3 and Ago4, but not to the negative control GAPDH (Fig. 4*B* and *C*). Therefore, the W2 and W3 peptides can interact with the four human Ago proteins. Interestingly, the binding to these Ago proteins was lost in the presence of the F2 peptide (Fig. 4*B*). Collectively, these data support a role for the W283 residue in Ago-binding *in vitro*. Contrary to the F2 peptide, the F3 mutant peptides remained competent in interacting with the four Ago proteins (Fig. 4*B*). To further investigate whether the W293 residue is functional *in vitro*, we decided to synthesize a peptide carrying a substitution of the tryptophan residue into alanine (Fig. 4*A*), a point mutation that has been the most extensively used on W-rich proteins to alter Ago-binding (19). We found that the resulting A3 peptide no longer interacted with Ago proteins (Fig. 4*C*), supporting a role for the W293 motif in Ago-binding *in vitro*. Altogether, these data indicate that both the W283 and W293 motifs behave as Ago-binding motifs *in vitro*. Nevertheless, structural modeling of LegK1 revealed that the W293 residue is buried into the kinase domain and thus unlikely accessible for interaction at the steady state (*SI Appendix*, Fig. S2*B*). Instead, we observed that the W283 is covered by a small structural element but could potentially be accessible upon conformational changes at the level of this segment. Collectively, these data suggest that LegK1 might interact with Ago proteins in a W-dependent manner, most likely through conformational changes (see discussion section).

**Fig. 4.**
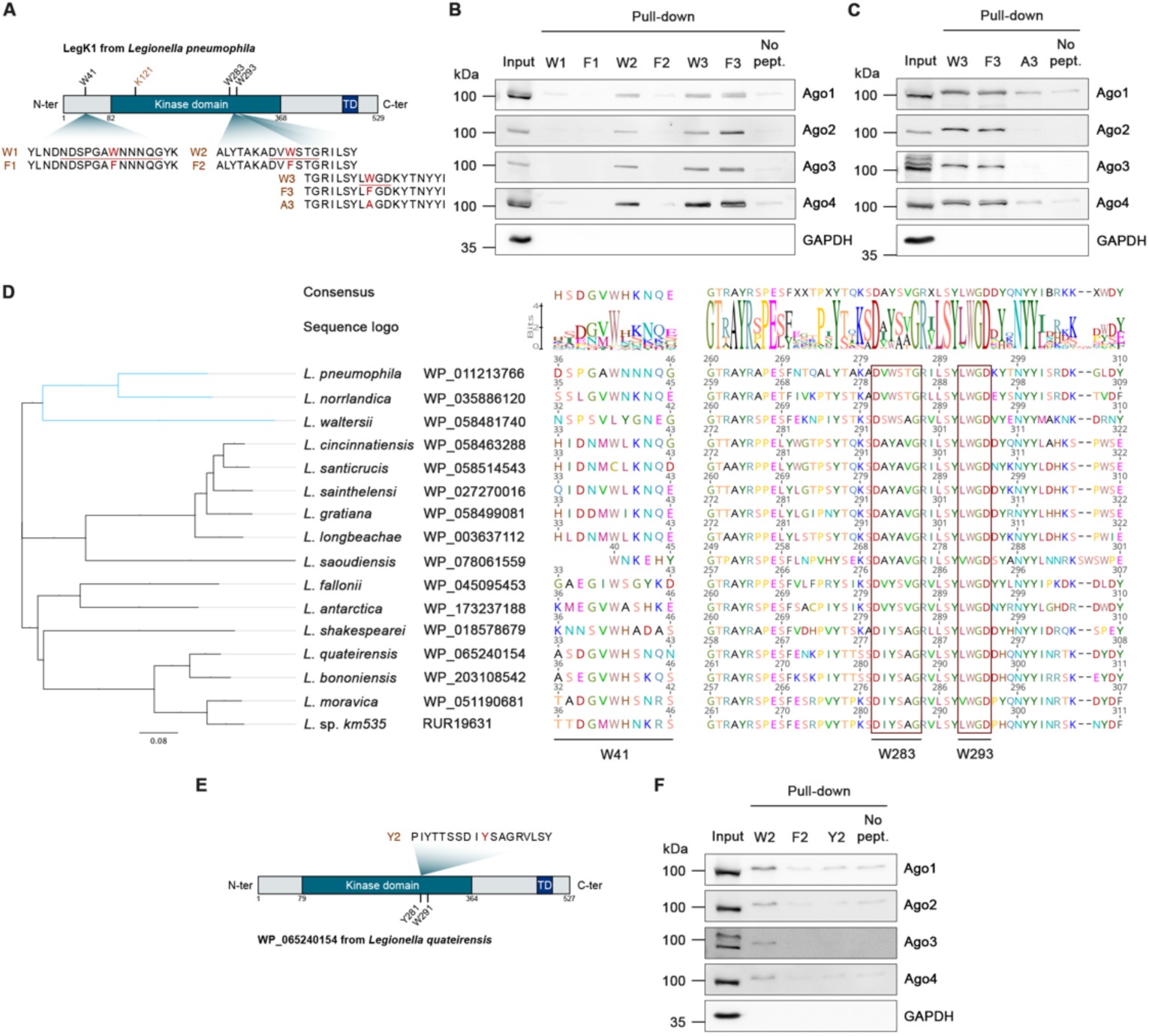
The W283 and W293 motifs of LegK1 bind to human Ago proteins and exhibit sequence conservation among the homologs of *Legionella*. (*A*) Schematic representation of LegK1 from *L. pneumophila* along with the synthetic biotinylated peptides used for pull-down assays. Each peptide sequence containing the WT (W1, W2, W3) or mutated versions (F1, F2, F3, A3) of predicted W-motifs of LegK1 are depicted. (*B*) and (*C*) Immunoprecipitations of W-motif-containing peptides from HEK293T cells. Synthetic biotinylated peptides were mobilized on streptavidin beads, incubated with HEK293T cell lysates and further immunoprecipitated. Incubation of beads without peptides was used as negative control (no pept.). The presence of the human Ago proteins in cell lysate (input) and bound to the beads was assessed by immunoblotting using the indicated antibodies. (*D*) Alignment of LegK1 homologs from *Legionella* species around the three putative W-motifs. The protein sequence of LegK1 (*lpp1439* gene) from *Legionella pneumophila* strain Paris was used as a reference sequence to determine the presence of homologs in the *Legionellales* order (taxid: 445). An identity cutoff of 40%, an Expect (*E*)-value cutoff of 10^−5^ and a minimum percentage match length of subject and query of 65% were used. The set of homologous protein sequences was aligned using ClustalW2. (*E*) Schematic representation of LegK1 homologs from *L. quateirensis* along with the synthetic biotinylated peptide used for pull-down assays. (*F*) Immunoprecipitations of the peptides as depicted in (*B*) and (*C*), plus an additional peptide (Y2), corresponding to the homologous W283-motif of *L. quatereinsis* from HEK293T cells, as depicted in (*B*) and (*C*). All data are representative of three independent experiments. GAPDH was used as negative control.

### The W283 and W293 motifs of *L*. *pneumophila* LegK1 exhibit sequence conservation among a subset or all *Legionella* species, respectively

To get some insights into the sequence conservation of the W-motifs of LegK1, we retrieved and aligned the protein sequences of putative LegK1 homologs from the *Legionella* protein sequences available on the National Center for Biotechnology Information (NCBI). More specifically, we selected for this analysis LegK1 homologous protein sequences exhibiting a protein identity of at least 40% with the *L. pneumophila* LegK1 strain Paris. According to these criteria, LegK1 was found present in 15 other *Legionella* species (2, 7) (Fig. 4*D*). Although the W41 motif was found conserved among all the *Legionella* species analysed, its surrounding residues were highly variable (Fig. 4*D*). Furthermore, they did not fulfill the typical features of residues neighbouring functional W-motifs, which are composed of small, polar and non-hydrophobic residues (43). This observation is thus consistent with our experimental data indicating that this Ago-binding motif is not functional (Fig. 4*B*). In contrast, the W293 motif of the strain *L. pneumophila* Paris, and its surrounding residues, were conserved among all *Legionella* species analysed (Fig. 4*D*). Therefore, if this W293 residue is functional, the equivalent W293 residues present in the different LegK1 homologous protein sequences analysed here must have maintained their Ago-binding capacity. When we analysed the sequence conservation of the W283 motif, and its surrounding residues, we found that these amino acids were also conserved among a cluster including *L. pneumophila*, *L. norrlandica* and *L. waltersii* (Fig. 4*D*, *blue clade*). However, we noticed sequence variations in the remaining *Legionella* species analysed, which were not only found in the surrounding residues, but also in the tryptophan itself (tryptophan to tyrosine residues (W>Y); Fig. 4*D*). Given that a tyrosine is one of the least probable residues that forms a functional W-motif (43), this observation suggested that the equivalent W283 motifs in the LegK1 protein sequences from these *Legionella sp.* had lost, or alternatively never gained, the ability to bind Ago proteins. Consistent with this hypothesis, a biotinylated peptide corresponding to the region of the *L. quateirensis* LegK1 sequence carrying the W>Y natural variation, named Y2, did not bind to any of the human Ago proteins (Fig. 4*E* and *F*). Therefore, the equivalent W283 motifs from LegK1 homologs should be functional in *L. norrlandica* and *L. waltersii,* but not in the remaining *Legionella* species.

### The kinase, Ago-binding and RNA silencing suppression activities of LegK1 are tightly interconnected

As the two functional W-motifs are embedded in the kinase domain of LegK1, we further investigated the relationship between the Ago-binding platform and the kinase activity of this effector. We examined the possible impact of single and combined W-motif mutations on the ability of LegK1 to phosphorylate IκBα, a known substrate of this effector protein (11). More specifically, we expressed the WT or the mutant versions of LegK1 in HEK293T cells, and further incubated the immunoprecipitates from each cell lysate with purified recombinant CBP-IκBα-His6. Western blot analyses from these cell-free extracts, using an antibody recognizing phosphorylation at Ser-32 of IκBα, revealed strong LegK1-induced phosphorylation of CBP-IκBα-His6 (Fig. 5*A*, *SI Appendix*, Fig. S6*A*). While the single LegK1-W283F mutant triggered comparable phosphorylation levels at Ser-32 of IκBα compared to the LegK1 WT version, the LegK1-W293F and LegK1-W293A mutants were partly and substantially compromised in this process, respectively (Fig. 5*A*). Importantly, the phosphorylation at Ser-32 of IκBα was fully abolished in the presence of LegK1-W283F-W293A, despite the stable accumulation of this double mutant in HEK293T cells. These effects were similar to the ones detected with the LegK1-3WF and the LegK1-KA mutants (*SI Appendix*, Fig. S6*A*). Therefore, the combined mutations in the two functional W-motifs of LegK1 are sufficient to abolish its catalytic activity.

**Fig. 5.**
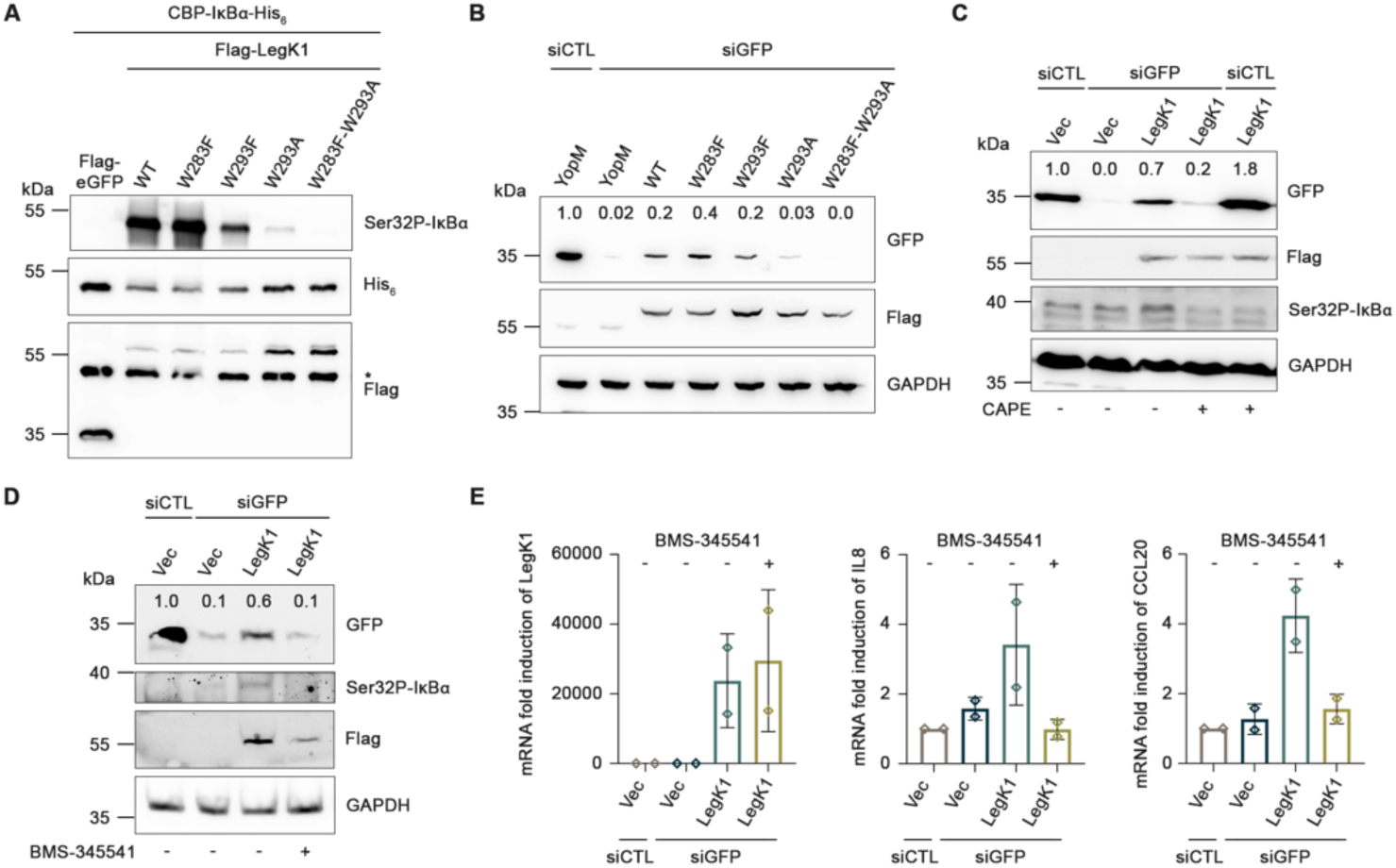
The kinase, RNA silencing suppression and NF-κB signaling activation activities of LegK1 are interconnected. (*A*) The LegK1-W293A and LegK1-W283F-W293A mutants are altered in their ability to phosphorylate IκBα. Kinase assay in the presence of LegK1 mutants. HEK293T cells were transfected with vectors expressing Flag-eGFP, 2xFlag-LegK1 WT, 2xFlag-LegK1-W283F, 2xFlag-LegK1-W293F, 2xFlag-LegK1-W293A and 2xFlag-LegK1-W283F-W293A. At 48h post-transfection, cells were lysed and subjected to a Flag immunoprecipitation. The resulting immunoprecipitates were incubated with purified CBP-IκBα-His6 recombinant proteins. The phosphorylation of the serine 32 of IκBα was analyzed by immunoblotting using a specific antibody. * represents aspecific bands. (*B*) The LegK1-W293A and -W283F-W293A mutants are altered in their ability to suppress silencing of the *GFP*-based siRNA reporter. *GFP*-based siRNA silencing reporter assay was performed in the presence of LegK1 mutant versions in HEK293T cells. Cells were co-transfected with pFlag-eGFP, negative control siRNA (siCTL) or GFP RNA duplexes, and vectors expressing Flag-YopM, 2xFlag-LegK1 WT, 2xFlag-LegK1-W283F, 2xFlag-LegK1-W293F, 2xFlag-LegK1-W293A or 2xFlag-LegK1-W283F-W293A for 48 hours. The eGFP protein levels are relative to the ones of GAPDH and further normalized to the YopM control condition (with siCTL), as depicted at the top of the GFP immunoblot. (*C*) and (*D*) The silencing suppression of the *GFP*-based siRNA reporter triggered by LegK1 is in part dependent on its ability to activate NF-κB signaling. *GFP*-based siRNA silencing reporter assay was performed in the presence of LegK1 in HEK293T cells treated with CAPE (*C*) or BMS-345541 (*D*). The experiment was performed as described in (*B*). Cells were treated with DMSO or 25 μg/mL of CAPE for 1h or 5 μM of BMS-345541 for 24h. The phosphorylation of the serine 32 of IκBα was used as a marker of NF-κB signaling activation. The eGFP protein levels are relative to the ones of GAPDH and further normalized to the empty vector (Vec) control condition (with siCTL), as indicated at the top of the GFP immunoblot. (*E*) BMS-345541 dampens the LegK1-triggered induction of *IL8* and *CCL20*. RT-qPCR analyses depicting the mRNA fold induction of *LegK1* (left panel), *IL8* (middle panel) and *CCL20* (right panel). The results shown are representative of two (*A*) and (*E*) or three (*B*), (*C*), (*D*) independent experiments.

We next assessed the possible impact of these W-motif mutations on the silencing suppression activity of LegK1 using the siRNA-based *GFP* reporter in HEK293T cells. We found that the LegK1-W293A mutant, and particularly the LegK1-W283F-W293A double mutant, were altered in their ability to suppress siRNA-guided silencing, as revealed by an impaired GFP derepression (Fig. 5*B*). These effects were comparable to the ones observed in HEK293T cells expressing YopM, LegK1-3WF or LegK1-KA mutants (Fig. 1*E*). We conclude that the W293 motif has a major role in LegK1-triggered silencing suppression and that both the W283 and W293 motifs might cooperatively orchestrate this process. Altogether, these data suggest that the kinase, Ago-binding and RNA silencing suppression activities of LegK1 are tightly interconnected. Nevertheless, we cannot exclude that the W293A substitution, which appears to trigger a destabilization effect *in silico* (*SI Appendix*, Fig. S2*A*), would alternatively, or additionally, alter the folding of the LegK1 protein, thereby impacting its kinase activity and consequently its ability to suppress RNA silencing.

### LegK1-induced NF-κB signaling contributes to RNA silencing suppression

Given that the kinase and silencing suppression activities of LegK1 were found interconnected, we reasoned that LegK1 could directly phosphorylate human Ago proteins to alter their functions. To test this hypothesis, we investigated whether LegK1 could phosphorylate human Ago2. However, we did not observe any increase in the phosphorylation status of the human Ago2 protein in the presence of LegK1 (*SI Appendix*, Fig. S6*B*-*G* and *Supporting Information Text*), indicating that LegK1 is unlikely to phosphorylate this host factor. Instead, these results suggested that LegK1 must suppress RNA silencing by phosphorylating other RISC component(s) and/or proteins that indirectly regulate RNA silencing activity. Because LegK1 has been shown to directly phosphorylate IκBα, resulting in a potent NF-κB signaling activation (11), and because its catalytic activity was found here to be essential for RNA silencing suppression (Fig. 1*E*-*G*), we further investigated whether LegK1-triggered NF-κB signaling activation could contribute to silencing suppression. For this purpose, we tested if the Caffeic Acid Phenethyl Ester (CAPE), a well-established NF-κB signaling inhibitor (49), could alter the ability of LegK1 to suppress silencing of the siRNA-based *GFP* reporter in HEK293T cells. Interestingly, while LegK1 triggered a significant derepression of siRNA-directed silencing of the *GFP* reporter, compared to the control vector, we found that this effect was significantly reduced upon CAPE treatment, which concomitantly prevented LegK1-induced phosphorylation at Ser-32 of IκBα (Fig. 5*C*). Of note, the low GFP protein accumulation detected in this condition was not due to a destabilization of the GFP in response to CAPE, because a normal GFP protein level was observed upon CAPE treatment of HEK293T cells expressing LegK1 and control siRNAs (Fig. 5*C*). The CAPE-dependent suppression of NF-κB signaling can thus restore the ability of siRNAs to silence the *GFP* reporter in the presence of LegK1. Because CAPE is known to exhibit broad molecular effects, beyond its known inhibitory effect on NF-κB signaling, we decided to conduct the same assay in the presence of a more selective NF-κB signaling inhibitor. The 4(2’-aminoethyl)amino-1,8-dimethylimidazo(1,2-a)quinoxaline, also known as BMS-345541, inhibits the catalytic subunits of IKK but does not alter the activity of 15 other tested kinases (50). Likewise, BMS-345541 does not affect c-Jun and STAT3 phosphorylation, as well as mitogen-activated protein kinase-activated protein kinase 2 activation in human cells. We found that BMS-345541 not only suppresses the ability of LegK1 to induce the NF-κB-dependent marker genes *interleukine 8* (*IL-8*) and *C-C motif chemokine ligand 20* (*CCL20*), but also to derepress the siRNA-based *GFP* reporter system (Fig. 5*D* and *E*). Our data, with two well-characterized NF-κB signaling inhibitors, support a link between LegK1-mediated NF-κB signaling activation and RNA silencing suppression. Altogether, our results demonstrate that the silencing suppression activity of LegK1 is in part achieved through the activation of NF-κB signaling directed by its kinase activity.

### LegK1 promotes the growth of *L. pneumophila* in both *Acanthamoeba castellanii* and human macrophages

To determine whether LegK1 contributes to the virulence of *L. pneumophila*, we next infected *Acanthamoeba castellanii,* a natural host of *L. pneumophila*, with the *Lpp* WT and Δ*legK1* strains and further monitored bacterial numbers during the course of the infection. Interestingly, the growth of the *Lpp* Δ*legK1* strain was reduced in *A. castellanii* compared to that of *Lpp* WT (Fig. 6*A*). Because *L. pneumophila* can accidentally infect humans where it replicates in alveolar macrophages, we conducted the same assay in macrophages derived from THP-1 cells. Similarly, a reduced intracellular replication of the Δ*legK1* strain was observed compared to the WT strain (Fig. 6*B*). These growth defects were partially complemented in both *A. castellanii* and human macrophages, when LegK1 was expressed *in trans* in the *Lpp* Δ*legK1* strain (Fig. 6*C*, *SI Appendix*, Fig. S7*A*). These data indicate that the reduced growth of *Lpp* Δ*legK1* in both *A. castellanii* and human macrophages is due to the deletion of *legK1*. They also provide evidence that LegK1 is a *bona fide* virulence factor of *L. pneumophila* that promotes bacterial growth in both amoeba and human macrophages.

**Fig. 6.**
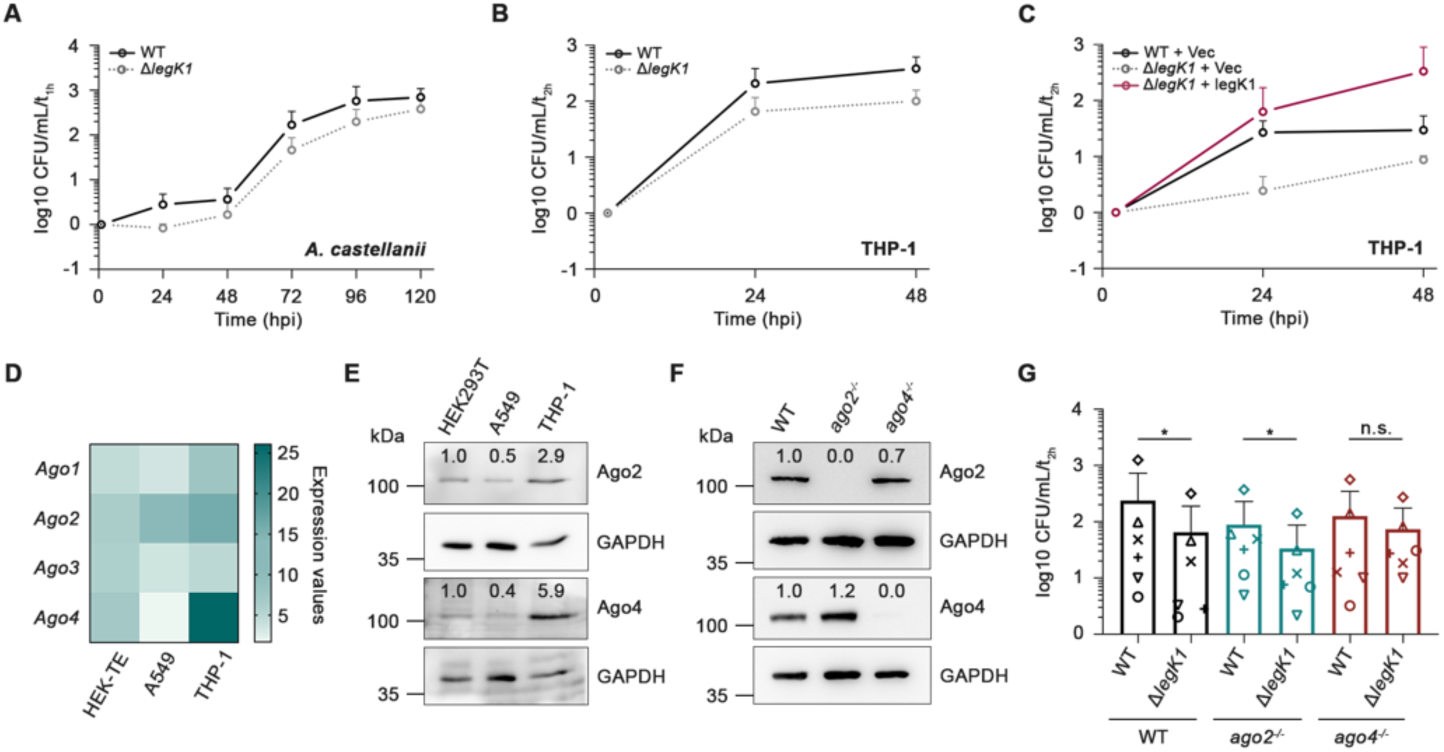
LegK1 contributes to RNA silencing suppression during infection and promotes bacterial growth by genetically targeting human Ago4. (*A*) LegK1 promotes *Lpp* growth in *A. castellanii*. *A. castellanii* were infected with *Lpp* WT and Δ*legK1* mutant strains at a MOI of 0.1 at 20°C. Intracellular replication was determined by recording the number of colony-forming units (CFU) by plating on buffered charcoal yeast extract (BCYE) agar. Results are shown as log10 ratio CFU/mL, where CFUs were normalized with the associated condition at 1h post-infection, corresponding to the entry of bacteria in host cells. (*B*) LegK1 promotes *Lpp* growth in human macrophages. THP-1 monocyte-derived macrophages were infected with *Lpp* WT or Δ*legK1* mutant strains at a MOI of 10 at 37°C. Results are shown as log10 ratio CFU/mL, where CFU were normalized with the associated condition at 2h post-infection, corresponding to the entry of bacteria in host cells. (*C*) Complementation assays in THP-1 cells. Cells were infected at a MOI of 10 at 37°C with the *Lpp* WT and Δ*legK1* mutant strains carrying the empty vector pBC-KS (Vec) or the complementing plasmid pBC-KS-LegK1. Intracellular growth was determined as in (*B*). (*D*) Heatmap of the four Ago transcript levels in HEK-TE, A549 and THP-1 cell lines. The RNA-sequencing data were recovered from the Cancer Cell Line Encyclopedia (Primary ID: E-MTAB-2770) and are reported as linear values. (*E*) Immunoblot of Ago2 and Ago4 in HEK293T, A549 and THP-1 cell lines. (*F*) Immunoblot of Ago2 and Ago4 in *ago2*^-/-^ and *ago4*^-/-^ CRISPR-Cas9-based knockout THP-1 cell lines. GAPDH was used as a loading control. (*G*) The growth defect of the *Lpp* Δ*legK1* strain is rescued in human macrophages depleted of Ago4. WT*, ago2 ^-/-^* and *ago4 ^-/-^* THP-1 monocyte-derived macrophages were infected with *Lpp* WT or Δ*legK1* strains at a MOI of 10. Bacterial titers were monitored at 24 hours post-infection (*hpi*) and were analysed as described in (*B*). Data are represented as ± SD. Data are representative of three (*A*, *E* and *F*), or six (*B, C* and *G*) independent experiments. Plotted data of infected THP-1 WT cells in (*C*) are the same as the data shown in (*G*) at 24 hpi. Linear regression analyses were performed in (*A*-*C*), highlighting significant growth differences between cells infected with WT or Δ*legK1* + LegK1 and Δ*legK1* strains. Statistical significance was determined using two-sided Wilcoxon rank-sum test in (*G*), *, P< 0.05; n.s., not significant.

### Human Ago4 is a functionally relevant genetic target of LegK1 in infected-macrophages

We next analysed whether LegK1 genetically targets human Ago functions to promote *L. pneumophila* virulence. Using available RNA-seq datasets (51), we first noticed that human *Ago2* and *Ago4* mRNAs were the most abundant *Ago* transcripts in THP-1 cells (Fig. 6*D*). This analysis supports recent findings showing that *Ago4* is highly expressed in immune-related cells (33). Consistently, we detected a higher Ago4 protein accumulation in THP-1 cells compared to HEK293T and A549 epithelial cells, which was also detected for Ago2, albeit to a lesser extent (Fig. 6*E*). Based on these data, we decided to knock-out human Ago2 or Ago4 in THP-1 cells for further functional assays. A single guide RNA approach was used to generate CRISPR/Cas9-based deletions in the second exon of the Ago2 and in the third exon of the Ago4 loci. The resulting pools of *ago2*^-/-^ and *ago4*^-/-^ THP-1 lines were found respectively deprived of Ago2 and Ago4 proteins compared to THP-1 control cells (Fig. 6*F*). Macrophages derived from the *ago2*^-/-^ and *ago4*^-/-^ THP-1 cells were infected with the *L. pneumophila* WT or Δ*legK1* strains and bacterial numbers were monitored at 24 hpi. A normal growth of the WT strain was achieved in *ago2*^-/-^ and *ago4*^-/-^ cells compared to control cells (*SI Appendix*, Fig. S7*B*), indicating that neither human Ago2 nor Ago4 control the growth of this bacterium in those conditions. Importantly, however, the growth defect of the *Lpp* Δ*legK1* strain was partially rescued in *ago4*^-/-^, but not in *ago2*^-/-^, macrophages (Fig. 6*G*). This indicates that the lack of Ago4 facilitates the growth of this bacterial mutant in macrophages. Therefore, this rescue experiment demonstrates that human Ago4 is a functionally relevant genetic target of LegK1 in infected-macrophages.

## Discussion

Here, we show that the type IV-secreted protein LegK1 from *L. pneumophila* can efficiently suppress RNA silencing in human cells. More specifically, LegK1 dampened the silencing of the Ago2-dependent siRNA-based *CXCR4* reporter (*SI Appendix*, Fig. S1). It also suppressed the silencing of a siRNA-based *GFP* reporter system, which depends, at least in part, on human Ago1 and Ago2 (*SI Appendix*, Fig. S1). In addition, we found that LegK1 was able to efficiently suppress a let-7a miRNA-based *luciferase* reporter with bulged target site (Fig. 1). Importantly, these silencing suppression effects relied on a predicted W-dependent Ago-binding platform, which contributed to the interaction of LegK1 with human Ago proteins (Fig. 1 and 2). Thus, these data indicate that LegK1 acts as a *bona fide* BSR by efficiently suppressing both siRNA- and miRNA-activities in human cells, presumably through its Ago-binding platform. The use of such Ago-binding platform is therefore unlikely restricted to VSRs, as previously thought (39, 42), but can be potentially exploited by at least one secreted virulence protein from a human pathogenic bacterium.

LegK1 was not only able to bind human Ago1, Ago2 and Ago4 through its W-motifs, but was also recovered in protein complexes containing TNRC6A, PABPC1 and DDX6 (Fig. 2). Because these factors are part of the HMW-miRISC (20), our data suggest that LegK1 interferes with the function of mature RISCs that are engaged in the silencing of miRNA targets. Intriguingly, we also noticed that the WS and LWG residues, which are present in the C-terminal domain of TNRC6 proteins, and known to be required for the direct interaction with components of deadenylase complexes (52–55), are also part of the W283 (DV<u>WS</u>TG) and the W293 (<u>LWG</u>D) motifs of LegK1. These observations suggest that LegK1 might additionally use its W-motifs to physically interact with deadenylase complexes. This hypothesis is consistent with our co-immunoprecipitation data showing that LegK1 interacts with DDX6, a direct partner of CNOT1 (52, 56). Based on these data, we propose that LegK1 might trigger silencing suppression by altering the function of HMW-RISCs. Such a mode of action would be well-adapted to rapidly repress RNA targets that fine-tune innate immune responses during *L. pneumophila* infections (35). It would also be analogous to the mode of action of the *Sweet potato mild mottle virus* (SPMMV) VSR P1, which was shown to interact with *Nicotiana benthamiana* AGO1 through W-motifs and to co-fractionate with RISC containing AGO1-loaded small RNAs (42).

We also found that the two W-motifs from the kinase domain of LegK1, namely W283 and W293, likely contribute to the binding of LegK1 to human Ago proteins (Fig. 4). Interestingly, these tryptophan residues were found interspaced by 9 amino acids, which is a typical feature of functional W-motifs embedded in the ABD of TNRC6 proteins, which are in most cases separated by 8 to 14 amino acids from each other (17, 19). This amino acid organization is thought to form a flexible linker that facilitates the interaction of W-motifs with the three W-binding pockets exposed at the surface of human Ago2 and Ago4 proteins, which are separated by a distance of 25 Å from each other (19). Based on these findings, we propose that the W283 and W293 motifs of LegK1 might cooperate with each other, upon possible conformational changes of the LegK1-Ago protein complexes (see following section), to ensure a tight association between this bacterial protein and the W-binding pockets of human Ago proteins.

To get further insights into the functional relevance of the W283 and W293 motifs of LegK1 in Ago-binding, we analyzed their sequence conservation across *Legionella* species. We found that the region corresponding to the *L. pneumophila* W293 motif, and its surrounding residues, are conserved among several *Legionella* species. The region corresponding to the W283 motif was also found conserved in *L. norrlandica* and *L. waltersii* species. In contrast, amino acid sequence variations were observed in the tryptophan and surrounding residues of the LegK1 protein sequences derived from the remaining analysed *Legionella* species. Furthermore, they prevented Ago-binding of a peptide derived from the *L. quateirensis* LegK1 protein sequence (Fig. 4). Therefore, the conservation of the W283 and W293 motifs, observed in some *Legionella* species, supports the hypothesis that these W-motifs cooperate between each other to ensure tight binding to Ago proteins. Nevertheless, our structural modeling also indicated that the W293 residue is buried into the kinase domain of LegK1 and thus is unlikely to be accessible for interactions at the steady state (*SI Appendix*, Fig. S2*B*). Therefore, we cannot exclude the possibility that the substitutions in W293 alternatively, or additionally, alter the LegK1 conformation, thereby impacting its ability to interact with Ago proteins. Instead, we observed that the W283 residue would be potentially accessible upon conformational changes (*SI Appendix*, Fig. S2*B*). These observations suggest that LegK1 interacts with Ago proteins in a W-independent manner, and/or in a W-dependent manner, but through conformational changes, which would possibly occur through protein partners from the miRISC and/or post-translational modifications. This would in turn render, the W-motif(s) accessible for interaction with the W-binding pockets of Ago PIWI domains. Unravelling the structure and dynamics of LegK1-Ago protein complexes, from human and/or protist cells, will be necessary to test these hypotheses in future studies.

By disabling the two functional W-motifs from the Ago-binding platform of LegK1, we found that this effector was fully compromised in its ability to phosphorylate IκBα and to suppress RNA silencing (Fig. 5). These data are consistent with the central role played by the kinase activity of LegK1 in RNA silencing suppression (Fig. 1), and suggest a tight interconnection between the kinase and Ago-binding activities of LegK1. In addition, by using two distinct and well-characterized NF-κB signaling inhibitors, we discovered that the blockage of NF-κB signaling activation in human cells expressing LegK1 reduced the siRNA-directed silencing of the *GFP* reporter system (Fig. 5). These data unveil an unexpected cross-talk between LegK1-mediated activation of NF-κB signaling and silencing suppression. They also highlight an interconnection between the kinase, RNA silencing suppression and NF-κB signaling activation activities of LegK1, which remains to be characterized in more depth in the future. It will notably be important to assess if cellular pools of human Ago proteins could be recovered in protein complexes containing NF-κB-related factors, and whether LegK1 acts on specific components of such complexes, besides IκBα and Ago proteins, to suppress RNA silencing while enhancing NF-κB signaling.

The long-lasting coevolution of *Legionella* in a wide range of amoebae and ciliated protozoa has shaped the *Legionella* genomes (6, 7). The presence of a large repertoire of eukaryotic-like proteins in the *Legionella* genomes, and their phylogenetic analyses, indicate that these proteins must have been acquired through horizontal gene transfer from protist host organisms (7, 9, 10, 57). Most of the virulence strategies used by *L. pneumophila* are required to replicate in both protists and human macrophages (2, 3). Consistently, we showed that the *Lpp* Δ*legK1* strain was altered in its ability to replicate in both *A. castellanii* and human macrophages (Fig. 6), supporting a physiologically relevant role of LegK1 in *L. pneumophila* virulence. In addition, we found that the growth defect of the *Lpp* Δ*legK1* strain was partially rescued in Ago4-deficient human macrophages (Fig. 6). This finding suggests that the targeting of Ago proteins by LegK1 is likely a major virulence function of this effector, which might also be required to promote growth of *L. pneumophila* in its natural hosts. This assumption is supported by the fact that amoebae genomes like *Dictyostelium discoideum*, possess canonical Ago proteins, some of which being even experimentally characterized in RNA silencing (58). Here, by searching for the presence of Ago-like factors in the recently released genome sequence of the *A*. *castellanii* strain used in this study (59), we recovered two candidate Ago-like proteins containing canonical PIWI and PAZ domains (*SI Appendix*, Fig. S7*C*), which is consistent with a recent report (58). Importantly, this observation is not restricted to *A*. *castellanii* as canonical Ago proteins were retrieved in available genomes from several Amoebozoa species (58), which are all possible natural hosts of *L*. *pneumophila*. In contrast, none of the NF-κB-related domains were identified in the genomes of amoebae, including the *A. castellanii* strain used herein (3, 59). Based on the lack of NF-κB-related genes in amoebae genomes, and on the presence of canonical Ago proteins in these natural hosts, we propose that the ability of LegK1 to suppress Ago activities is likely the primary virulence function of this bacterial effector.

Antiviral RNA interference (RNAi) has been extensively characterized in plants, fungi, insects and worms (60). In mammals, emerging evidence indicates that antiviral RNAi also operates in embryonic stem cells, adult stem cells and progenitor cells, in which the antiviral interferon (IFN) response –known to repress RNAi– is less active (61–64). In these tissues/cell types, canonical antiviral sRNAs readily accumulate during viral infections and inhibit viral replication in an Ago2-dependent manner (63). Interestingly, mouse Ago4 was also recently shown to restrict the replication of different viruses in IFN-competent cells, including *influenza A virus* (IAV) (33). Importantly, mouse Ago4 was also shown to dampen IAV replication *in vivo*, further supporting a key role of this factor in antiviral resistance (33). In the present study, we first showed that Ago4 – and Ago2 to a lesser extent– were the most expressed Ago genes in immune-related cells (Fig. 6). Significantly, although a normal growth of the *Lpp* WT strain was observed in human *ago2^-/-^* and *ago4^-/-^* macrophages, we found that the growth defect of the *Lpp* Δ*legK1* mutant was partially rescued in Ago4-deficient cells (Fig. 6). This provides genetic evidence that LegK1 must suppress the function of Ago4 to promote growth of *L. pneumophila* in IFN-competent macrophages. However, future investigations will be necessary to determine whether mammalian Ago4 could orchestrate antibacterial immune responses, beyond its known antiviral function.

In conclusion, this study reports on the first bacterial effector that directly suppresses RNA silencing by physically interacting with Ago proteins, in part through W-motifs. Intriguingly, we have also identified W-motifs in an appropriate sequence context for Ago-binding in effectors from various human pathogenic bacteria, fungi and parasites (*SI Appendix*, Fig. S8). These observations suggest that a wide range of non-viral mammalian pathogens and parasites might have evolved an analogous mode of action to promote virulence. It will thus be interesting to establish if the discoveries made on LegK1 hold true for other virulence/parasitic factors from non-viral mammalian pathogens/parasites. It will also be worth developing innovative strategies to counteract this potentially widespread virulence mechanism in order to control infectious and parasitic diseases.

## Materials and Methods

Please see *SI Appendix* for a detailed description of the *Materials and Methods*.

### Human cell lines and *Acanthamoeba castellanii* culture

HeLa cells (ATCC^®^ CCL-2™), control (CTL), *ago1*^-/-^, *ago2*^-/-^, *ago1*^-/-^/*ago2*^-/-^ and *dicer*^-/-^ CRISPR/Cas9-based HeLa cell lines (65) (this study), Human Embryonic Kidney 293T (HEK293T) cells (ATCC^®^ CRL-3216™) and HEK293 cells stably expressing the macrophage Fcɣ-RII receptor, gift from Craig Roy (48), were cultured in high glucose Dulbecco’s modified Eagle’s medium (DMEM) (containing 4.5 g/l of glucose) (Dominique Dutscher, L0103-500). A549 cells (ATCC^®^ CCL-185) were cultured in Ham’s F-12K Nutrient Mixture (Kaighn’s Modification) (Corning, 10-025-CVR). THP-1 (ATCC^®^ TIB-202™) and THP-1 *ago2*^-/-^ and *ago4*^-/-^ cell pools (Synthego®, this study) were cultured in Gibco Roswell Park Memorial Institute (RPMI) 1640 medium (Dominique Dutscher, L0498-500). All media were supplemented with 10% fetal bovine serum. Cells were maintained at 37°C in a humidified 5% CO2 atmosphere. In addition, all human cell lines were regularly tested negative for mycoplasma contamination. *Acanthamoeba castellanii* (ATCC^®^ 50739) was cultured in PYG 712 medium (2% proteose peptone, 0.1% yeast extract, 0.1 M glucose, 4 mM MgSO4, 0.4 M CaCl2, 0.1% sodium citrate dihydrate, 0.05 mM Fe (NH4)2(SO4)2 x 6H2O, 2.5 mM NaH2PO3, 2.5 mM K2HPO3) at 20°C for 72 h prior to harvesting for *L. pneumophila* infection.

### *GFP*-based silencing reporter system analyses

The suppression activities on siRNA-guided *GFP* silencing reporter were determined by Western blot analyses. Cell lines were seeded in 24-well plates and co-transfected for 48h with 200 ng of pFlag-eGFP and 30 pmol of GFP RNA duplex (RNA GFP duplex I, Thermoscientific, P-002048-01-20) or AllStars Negative Control siRNA, and with 1 μg of p2xFlag-LegK1-WT, -KA, -3WF, -W283F, -W293F, -W293A, -W283F-W293A, pFlag-YopM or empty vector as negative control, or pFlag-T6B as positive control using Lipofectamine 2000. Transfected cells were washed with 1X PBS and lysed in 100 μL of 1X Laemmli Loading Buffer. For each condition, 40 μL of samples were loaded on SDS-PAGE and subjected to Western Blot analysis. In some experiments, cells were transiently co-transfected with siRNA-guided silencing reporters and p2xFlag-LegK1 or pFlag empty control plasmid or pFlag-YopM, to ensure that each transfection received the same amount of total DNA. For the inhibition of NF-κB signaling, cells were then treated for 1h with 25 μg/mL of Caffeic Acid Phenethyl Ester inhibitor (CAPE, Santa Cruz Biotechnology, sc-200800), BMS-345541 (Sigma, B9935) or DMSO. Cell extracts were subjected to Western blot analysis, as described previously. Quantification of eGFP protein from Western blot analysis was performed by densitometric analysis, using the Fiji (ImageJ) software, and was then normalised to the YopM-siCTL or empty vector-siCTL conditions.

### *A. castellanii* and THP-1 infection assays

*A. castellanii* were washed once with Infection Buffer (PYG 712 medium without proteose peptone, glucose and yeast extract) and seeded in 25 cm^2^ flasks at a concentration of 4×10^6^ cells per flask. *L. pneumophila* WT and mutant strains were grown on BCYE agar to stationary phase, diluted in Infection Buffer and mixed with *A. castellanii* at a Multiplicity Of Infection (MOI) of 0.1 or 1 for complementation experiments. The infection was performed at 20°C. After a 1h-invasion period, the *A. castellanii* layer was washed three times with Infection Buffer. For every time point, the intracellular multiplication of bacteria was monitored collecting 300 µL of sample, that were centrifuged (14,500 rpm, 10 min), vortexed to break up amoeba, and plated on BCYE medium. For THP-1 infection, cells were seeded into 12-well tissue culture trays at a density of 2×10^5^ cells per well and pretreated 72 h with 10^-8^ M phorbol 12-myristate 13-acetate (PMA) (Sigma) in 5% CO2 at 37°C, to induce differentiation into macrophage-like adherent cells. Stationary phase *Legionella* were resuspended in RPMI 1640 serum free medium and added to THP-1 cells monolayer at a MOI of 10. The infection was performed at 37°C. After a 2h-incubation, cells were washed three times with 1X PBS before incubation with serum-free medium. At 24h and 48h post-infection, THP-1 cells were lysed with 1X PBS-0.1% Triton X-100. The *Legionella* titers were monitored by counting the number of colony-forming units (CFU) after plating on BCYE agar.

## Data availability

All materials generated in this study are available upon request. Mass spectrometry data are available *via* ProteomeXchange with identifier PXD037279. Sequencing data corresponding to the mutant *DlegK1* have been submitted to Zenodo (https://zenodo.org/record/7190772). All data reported in this paper will be shared by the lead contact upon request.

## Acknowledgments

We thank Dr. Alice Lebreton, Dr. Julie Guignot, Dr. Benoit Garin, Dr. Lauriane Quenee, Dr. Clément Carré, Pr. Gunter Meister, Dr. Hervé Le Hir for providing the Staphylococcus aureus SH1000 genomic DNA, Pseudomonas aeruginosa PAO1 strain and Escherichia coli O157:H7 gDNA, Bordetella pertussis Tohama gDNA, Yersinia pestis gDNA, AutomiG reporter, a vector expressing T6B, pET22-6His-SUMO and pET28-CBP-6His, respectively. We thank Dr. Sophie Goudey for the generation of KO HeLa cell lines, and Jérôme Zervudacki for his help on Northern blot analyses. We also thank all members of the Navarro lab for their inputs, discussions and critical reading of the manuscript, as well as Dr. Alice Lebreton, Dr. Michael Mourez, and Miss Isabelle Barbosa for helpful advices and discussions. This work was supported by the European Research Council (281749, Silencing & Immunity, to L.N.), a grant from the European Molecular Biology Organization (EMBO) Young Investigator Program (to L.N.), an Action Thématique et Incitative sur Programme (ATIP)/Avenir Grant from the Fondation Bettencourt Schueller (FBS, to L.N.), an Emergence Grant from the Mairie de Paris (to L.N.), a CIFRE PhD program from the Association Nationale Recherche Technologie (ANRT, 2017/0009 to J.T), a fourth-year PhD program from the Fondation pour la Recherche Médicale (FRM, FDT202001010790 to J.T.) and a grant ANRS (ECTZ47170 to S.G.M.). Work in the C.B. laboratory was supported by the Institut Pasteur, the FRM grant N° EQU201903007847 and the Agence Nationale de la Recherche grant n°ANR-10-LABX-62-IBEID. We thank the group of Craig Roy for providing the HEK293-FcɣRII cells. We gratefully acknowledge the UtechS Photonic BioImaging, C2RT, Institut Pasteur, supported by the French National Research Agency (France BioImaging; ANR-10–INSB–04; Investments for the Future). PB acknowledges support from the LABEX DYNAMO (ANR-11-LABX-0011), and the LBT IT team for access to the LBT computing resources.

