## Supplementary material for "A *Legionella pneumophila* effector impedes host gene silencing to promote virulence": SI appendix

This manuscript was deposited as a preprint on bioRxiv, doi:
https://doi.org/10.1101/2022.11.16.516792

**This PDF file includes:**

SI Materials and Methods
Supporting text
Figures S1 to S9
Tables S1 to S3
SI References

### SI Materials and Methods

#### Human cell lines and culture

HeLa cells (ATCC® CCL-2™), control (CTL), *ago1*$^{-/-}$, *ago2*$^{-/-}$, *ago1*$^{-/-}$/*ago2*$^{-/-}$ and *dicer*$^{-/-}$ CRISPR/Cas9-based HeLa cell lines (1) (this study), Human Embryonic Kidney 293T (HEK293T) cells (ATCC® CRL-3216™) and HEK293 cells stably expressing the macrophage Fcɣ-RII receptor, gift from Craig Roy (2), were cultured in high glucose Dulbecco's modified Eagle's medium (DMEM) (containing 4.5 g/l of glucose) (Dominique Dutscher, L0103-500). CTL and *ago2*$^{-/-}$ CRISPR/Cas9-based HeLa cell lines were previously generated (1). LentiCRISPRv2-Ago1 and LentiCRISPRv2-Dicer expression plasmids expressing single-guide RNA targeting exon 3 of Ago1 (AAACTCATACACAGGCTTGCGATC) or Dicer (AAACCTGATCTGATAGGACAGCTC) genes respectively, were used to transfect HEK293T cells using Lipofectamine LTX (Life technologies, A12621). Forty-eight hours later, viral particles were harvested and used to transduce HeLa cells. Cells were selected in medium supplemented with puromycin for 15 days. Data acquisition and data analysis were performed on the Cochin Cytometry and Immunobiology core Facilities. Single cells were isolated from lentiCRISPRv2-Ago1 or lentiCRISPRv2-Dicer transduced cells using a BD Biosciences FACSAria III. Expression of Ago1 or Dicer in the cell clones was analyzed by SDS-PAGE and immunoblotting. One cell clone knockout was selected for further analysis. A549 cells (ATCC® CCL-185) were cultured in Ham's F-12K Nutrient Mixture (Kaighn's Modification) (Corning, 10-025-CVR). THP-1 (ATCC® TIB-202™) and THP-1 *ago2*$^{-/-}$ and *ago4*$^{-/-}$ cell pools (Synthego®, this study) were cultured in Gibco Roswell Park Memorial Institute (RPMI) 1640 medium (Dominique Dutscher, L0498-500). The percentages of editing efficiency after expansion were of 97% and 96% for THP-1 *ago2*$^{-/-}$ and *ago4*$^{-/-}$ cell pools, respectively. All media were supplemented with 10% fetal bovine serum. Cells were maintained at 37°C in a humidified 5% $CO_2$ atmosphere. In addition, all human cell lines were regularly tested negative for mycoplasma contamination.

#### *Acanthamoeba castellanii* culture

*Acanthamoeba castellanii* (ATCC® 50739) was cultured in PYG 712 medium (2% proteose peptone, 0.1% yeast extract, 0.1 M glucose, 4 mM $MgSO_4$, 0.4 M $CaCl_2$, 0.1% sodium citrate dihydrate, 0.05 mM Fe $(NH_4)_2(SO_4)_2$ x $6H_2O$, 2.5 mM $NaH_2PO_3$, 2.5 mM $K_2HPO_3$) at 20°C for 72 h prior to harvesting for *L. pneumophila* infection.

#### Bacterial strains and mutant constructions

*Legionella pneumophila* strain Paris WT (CIP 107629T) and *ΔlegK1* (*Δlpp1439*) mutant were cultured in N-(2-acetamido)-2-aminoethanesulfonic acid (ACES)-buffered yeast extract broth (BCY) or on ACES-buffered charcoal-yeast (BCYE) extract agar (3). When needed antibiotics were added for *L. pneumophila*: gentamycin 12.5 µg/ml, chloramphenicol 10 µg/ml. All strains were grown at 37°C. The *ΔlegK1* mutant was constructed by replacing the gene of interest with a gentamycin resistance cassette in the chromosome. The mutant allele was constructed using a three-step PCR technique (4) (*SI Appendix*, Table S1). Therefore, three overlapping fragments, corresponding to the upstream region, the antibiotic cassette and the downstream region of the gene of interest, were amplified independently and purified on agarose gel. The three resulting PCR products were mixed at the same molar concentration (15 nM) and a second PCR with flanking primer pairs was performed. The resulting PCR product, the gentamycin resistance cassette flanked by 500 bp regions homologous to *legK1* was introduced into *L. pneumophila* by natural competence for chromosomal recombination. Strains that had undergone allelic exchange were selected by plating on BCYE containing gentamycin and the mutant was verified by PCR and whole genome sequencing. Bacterial growth assays were performed by seeding the indicated strains in wells of a microtiter plate (TPP) containing 200µl of BCYE. The microtiter plate was incubated with shaking at 37°C for 48 hours in a Tecan Infinite multiwell plate reader (Tecan Group). Optical density measurements were taken at 600nm every 30 min. For complementation, the *L. pneumophila* WT and *ΔlegK1* mutant strains were transformed by electroporation (2.5kV, 200Ω and 25µF) with 100 ng of the empty plasmid or pBC-KS-LegK1. Transformants were selected by plating on BCYE containing chloramphenicol.

**Expression plasmids and constructions**

The expression vectors for bacterial candidates were generated using GATEWAY technology. Briefly, PCR amplification (*SI Appendix*, Table S1) of *Lysteriolysin O* and *α-Hemolysin* coding sequences was performed using pAD-hly-Myc plasmid and *Staphylococcus aureus* SH1000 genomic DNA as templates, respectively (gift from Dr. Alice Lebreton, IBENS, Paris, France). *BepB* coding sequence was amplified from *Bartonella Henselae* gDNA, *ExoY* from *Pseudomonas aeruginosa* PAO1 strain, *NleH1* from *Escherichia coli* O157:H7 gDNA (gift from Dr. Julie Guignot, Institut Cochin, Paris, France), *Pertussis toxin* from *Bordetella pertussis* Tohama gDNA (gift from Dr. Benoit Garin, Institut Pasteur, Paris, France), *YopM* from *Yersinia pestis* gDNA (gift from Dr. Lauriane Quenee, Institute Biosafety Officer at Caltech, Los Angeles, USA). Finally, the coding sequence of *legK1* (*lpp1439*) was PCR amplified using gDNA of the *Legionella pneumophila* strain Paris. All resulting PCR products were introduced into the pENTR/D-TOPO entry vector (Invitrogen, K240020). They were sequenced and then recombined into the GATEWAY binary destination vector pPURO-Flag-HA (gift from Dr. Sebastien Pfeffer, IBMC, Strasbourg, France), hereinafter referred to as pFlag, using LR clonase (Invitrogen, 11791020), allowing an expression under the control of the cytomegalovirus (CMV) immediate early promoter. The molecular characterization of LegK1 was then done in the HEK293T cell line, using a pPURO-2xFlag-HA vector, hereinafter referred to as p2xFlag. This plasmid was constructed from pFlag expression vector, by insertion of a second Flag using Gibson assembly (New England BioLabs, E5510S). The *legK1* coding sequence was then inserted from pENTR/D-TOPO-LegK1 into p2xFlag through the GATEWAY technology, as previously described. Point mutations were introduced in this plasmid to generate the kinase-dead mutant (LegK1-KA), the mutants on the three putative W-motifs (LegK1-3WF), on the two functional W-motifs (LegK1-W283F-W293A) or on the individual W-motifs (LegK1-W283F, -W293F, -W293A). All these point mutations were carried out using site-directed mutagenesis by PCR with appropriate primers containing the desired nucleotide changes (Supplementary Table 1), and followed by selection with DpnI digestion. A vector expressing TNRC6B-derived peptide (T6B) was used as positive control of the siRNA-guided GFP silencing reporter, and was kindly provided by Pr. Gunter Meister (5) (Universität Regensburg, Regensburg, Germany). The pGL3-CXCR4-2p and pRL-TK plasmids were a gift from Dr. Sébastien Pfeiffer. For siRNA-guided *GFP* silencing reporter, the plasmid pPURO-Flag-HA-eGFP was cloned in this study using GATEWAY technology, as previously described, and was hereinafter referred to as pFlag-eGFP. The let-7a luciferase reporter was constructed by cloning a single let-7a target site between the XhoI and NotI restriction sites from the multiple cloning site (MCS) located downstream of the renilla luciferase reporter in the psiCheck-2 vector, giving rise to the psiCheck-*let-7a* plasmid.

For immunofluorescence analyses, LegK1 WT and its derivatives forms (LegK1-KA and LegK1-3WF) were PCR amplified and inserted using XhoI-BamHI into pmCherry-C1 (Clonetech). Plasmids expressing human Ago2 and Ago4 fused to GFP in the N-terminal region of Ago2 and Ago4 were obtained from Addgene (11590 and 21536, respectively). Empty pmCherry-C1 and pEGFP-C1 (Clonetech) were used as negative controls. For recombinant protein purification, a DNA fragment encoding LegK1 amino acids 2 to 386 was amplified by PCR, digested with NotI and XhoI enzymes and inserted into pET22-$His_6$-SUMO plasmid (6), kindly given by Dr. Hervé Le Hir (IBENS, Paris, France). This construct allows the expression of $LegK1^{2–386}$ recombinant protein in fusion with an amino-terminal SUMO tag and a polyhistidine tag. The pET22-$His_6$-SUMO-$LegK1^{2-386}$-KA and pET22-$His_6$-SUMO-$LegK1^{2-386}$-3WF were generated using site-directed mutagenesis by PCR. The full-length GST-Ago2 was purified from the pGST-Ago2 plasmid, obtained from Addgene (24317). A construct including only the PIWI domain of Ago2 and a fraction of MID domain was generated by PCR amplification of Ago2 amino-acids 517 to 859. $PIWI^{517-859}$ DNA was digested by BamHI and HindIII enzymes and cloned into pET28-CBP-$His_6$, kindly given by Dr. Hervé Le Hir, allowing a fusion with the Calmodulin Binding Protein (CBP) and a polyhistidine tag in amino-terminal and in carboxyl-terminal ends, respectively. Finally, the full-length IκBα was purified from pET28-CBP-IκBα-$His_6$ plasmid. IκBα DNA was PCR amplified from pBABE-GFP-IκBα-wt (Addgene, 15263) and was digested and cloned into pET28-CBP-$His_6$, as described for pET28-CBP-$PIWI^{517-859}$-$His_6$. The negative control protein TAP, consisting of the CBP and $His_6$ tags, was purified from a pET28 plasmid derivative (kindly provided by Dr. Hervé Le Hir)[107]. For complementation assay, the full-length *legK1* gene was cloned under the control of its own promoter into pBC-KS (Stratagene,

212215) using SacI and KpnI restriction enzymes. All constructions were confirmed by Sanger sequencing.

**Human cell transfections**

For the detection of recombinant LegK1 WT or mutant proteins by Western blot analysis, HEK293T cells were seeded in 24-well plates at a density of $1.4x10^5$ cells per well pre-treated with poly-L-Lysine (Sigma, P8920), and were transiently transfected for 48h with either 500 ng of p2xFlag-LegK1-WT, -KA or -3WF plasmids using JetPrime (Polyplus, 114-15), according to the manufacturer's instructions.

For the characterization of the siRNA-based luciferase reporter, control (CTL), *ago1*$^{-/-}$, *ago2*$^{-/-}$, *ago1*$^{-/-}$/*ago2*$^{-/-}$ and *dicer*$^{-/-}$ HeLa cell lines were transfected with Lipofectamine 2000 (Invitrogen, 11668019). One day before transfection, HeLa cell lines were trypsinized, resuspended in DMEM, and seeded in 48-well plates at a density of $7x10^4$ cells per well. Cells were transiently co-transfected for 48h with 100 ng of pGL3-CXCR4-2p, 50 ng of pRL-TK as transfection control, and 250 pM of AllStars Negative Control siRNA (Qiagen, 1027280) or CXCR4 siRNA (GUUUUCACUCCAGCUAACACA, Eurofins Genomics). For the initial screening of bacterial effectors, WT HeLa cells were co-transfected as previously described, with an addition of 400 ng of vector expressing a bacterial candidate or HBx as positive control. Firefly and Renilla luciferase expressions were determined as described in the Dual-luciferase silencing reporter analyses section. For the siRNA-based GFP sensor, HeLa and HEK293T cell lines were seeded in 24-well plates at the same density as mentioned above. Cells were co-transfected for 48h with 200 ng of pFlag-eGFP and 30 pmol of GFP RNA duplex (RNA GFP duplex I, Thermoscientific, P-002048-01-20) or AllStars Negative Control siRNA, and with 1 μg of p2xFlag-LegK1-WT, -KA, -3WF, -W283F, -W293F, -W293A, -W283F-W293A, pFlag-YopM or empty vector as negative control, or pFlag-T6B as positive control using Lipofectamine 2000. GFP proteins levels were next analysed by Western blot. For the let-7a miRNA-based luciferase sensor, HEK293T cells were seeded in 24-well plates at the same density as previously described and cells were co-transfected with 200 ng of psiCheck-*let-7a* and 1 μg of plasmid expressing either WT or LegK1 mutants, YopM or T6B using Lipofectamine 2000 for 48h. GFP and Renilla protein levels were further analysed by Western blot analyses. For let-7a miRNA silencing reporter assay during infection, HEK293-FcγRII cells were seeded in 24-well plates at a density of $1x10^5$ cells per well pre-treated with poly-D-lysine (7) at 2 µg/ml and transiently transfected during 24h with 400 ng of psiCheck-*let-7a* using JetPrime.

For immunofluorescence analyses, HEK293-FcγRII cells were seeded in 12-well plates at a density of $5x10^4$ cells per well pre-treated with poly-D-lysine at 2 µg/ml and transiently transfected during 48h with 1µg of pEGFP and pmCherry plasmids using JetPrime.

For the co-immunoprecipitation assays, HEK293T cells were seeded in 10 $cm^2$ dishes at a density of $6.2x10^6$ cells and were transiently transfected during 48h with 10 μg of pFlag-eGFP or p2xFlag-LegK1-WT, -KA, -3WF using JetPrime. For the co-immunoprecipitation assay followed by RNAse A treatment (Thermo Scientific, R1253), HEK293T cells were seeded in the same manner and were co-transfected for 48h with 10 μg of pGFP-Ago2-WT and 10 μg of pFlag-T6B or 14 μg of p2xFlag-LegK1 using Lipofectamine 2000. To determine the catalytic activity of LegK1 mutants in the kinase assay, HEK293T cells were seeded in 6 $cm^2$ dishes at a density of $8.4x10^5$ cells, and were transiently transfected for 48h with 4 μg of pFlag-eGFP or p2xFlag-LegK1-WT or p2xFlag-LegK1 mutants using JetPrime. Finally, for the mass spectrometry analysis, HeLa cells were seeded in 15 $cm^2$ dishes at a density of $1.4x10^7$ cells and were transfected for 24h with 20 μg of pFlag-eGFP, p2xFlag-LegK1-WT or -KA using JetPrime.

For RNA extraction, HEK293T cells were seeded in 12-well plate at a density of $2.8x10^5$ cells per well, and were transiently transfected for 48h with 800 ng of pFlag-eGFP or 1 μg of p2xFlag-LegK1-WT using JetPrime.

#### *A. castellanii* and THP-1 infection assays

*A. castellanii* were washed once with Infection Buffer (PYG 712 medium without proteose peptone, glucose and yeast extract) and seeded in 25 $cm^2$ flasks at a concentration of $4x10^6$ cells per flask. *L. pneumophila* WT and mutant strains were grown on BCYE agar to stationary phase, diluted in Infection Buffer and mixed with *A. castellanii* at a Multiplicity Of Infection (MOI) of 0.1 or 1 for complementation experiments. The infection was performed at 20°C. After a 1h-invasion period, the *A. castellanii* layer was washed three times with Infection Buffer. For every time point, the intracellular multiplication of bacteria was monitored collecting 300 µL of sample, that were centrifuged (14,500 rpm, 10 min), vortexed to break up amoeba, and plated on BCYE medium. For THP-1 infection, cells were seeded into 12-well tissue culture trays at a density of $2x10^5$ cells per well and pretreated 72 h with $10^{-8}$ M phorbol 12-myristate 13-acetate (PMA) (Sigma) in 5% $CO_2$ at 37°C, to induce differentiation into macrophage-like adherent cells. Stationary phase *Legionella* were resuspended in RPMI 1640 serum free medium and added to THP-1 cells monolayer at a MOI of 10. The infection was performed at 37°C. After a 2h-incubation, cells were washed three times with 1X PBS before incubation with serum-free medium. At 24h and 48h post-infection, THP-1 cells were lysed with 1X PBS-0.1% Triton X-100. The *Legionella* titers were monitored by counting the number of colony-forming units (CFU) after plating on BCYE agar.

#### Dual-luciferase silencing reporter analyses

The initial screening in HeLa cells was carried out using a siRNA-guided luciferase silencing reporter, consisting in the expression of the pGL3-CXCR4-2p vector (8), along with CXCR4 RNA duplexes. In parallel, a Renilla luciferase control reporter vector was used, corresponding to the pRL-TK. Transfected cells were washed with 1X PBS and then lysed in 65 µL of 1X Passive Lysis Buffer (Dual-Luciferase Reporter Assay System, Promega, E1910) by shaking for 15 min at room temperature. Firefly and Renilla activities were measured in 10 µL of cell lysate with the Luciferase reporter assay system, according to the supplier's protocol. Briefly, bioluminescence was initiated by automatic injection of 30 µL of Luciferase Assay Reagent II. After a 1 second-shaking and an additional 1 second-delay, Firefly luciferase emission signals were recorded on a TriStar LB 941 Multimode Microplate Reader luminometer (Berthold Technologies), using a 10 second-measurement period for each condition. The Renilla luciferase emission signals were then measured after injection of 30 µL of Stop & Glo Reagent, in the same manner as for Firefly luminescence detection. All measurements were carried out in three technical replicates for each independent experiment. Firefly luminescence expression was corrected with the Renilla control, the luminescence value of siCXCR4 condition was then normalised on siCTL condition, and further normalised by the negative control condition, corresponding to the CTL HeLa cell line or eGFP transfected WT HeLa cells.

The miRNA-guided luciferase silencing reporter consists in the expression of the psiCheck-*let-7a* vector composed of a non-targeted *firefly luciferase* which served as an internal reference control, and a Renilla luciferase construct carrying, downstream of its coding sequence, a let-7a complementary sequence containing mismatches opposite to nucleotides at positions 9, 10 and 11 of the let-7a sequence. Transfected HEK293T cells were prepared as described above for the siRNA-guided luciferase silencing reporter. For let-7a miRNA silencing reporter assay during infection, HEK293-FcγRII cells were transfected for 24h and then were infected with *L. pneumophila* WT strain or its derivative mutants grown until post-exponential phase (OD~4). Before cell infection, the bacteria were pre-opsonized by incubating them with an anti-FlaA antibody for 30 minutes at 37˚C. Then, the cells were infected with a MOI of 10. At the indicated time points, cells were washed with 1X PBS and then lysed in 130 µL of 1X Passive Lysis Buffer (Dual-Luciferase Reporter Assay System, Promega, E1910) by shaking for 15 min at room temperature.

#### *GFP*-based silencing reporter system analyses

The suppression activities on siRNA-guided *GFP* silencing reporter were determined by Western blot analyses. Transfected cells were washed with 1X PBS and lysed in 100 µL of 1X Laemmli Loading Buffer. For each condition, 40 µL of samples were loaded on SDS-PAGE and subjected to Western Blot analysis. In some experiments, cells were transiently co-transfected with siRNA-guided silencing reporters and p2xFlag-LegK1 or pFlag empty control plasmid or pFlag-YopM, to ensure that each transfection received the same amount of total DNA. For the inhibition of NF-κB signaling, cells were then treated for 1h with 25 μg/mL of Caffeic Acid Phenethyl Ester inhibitor (CAPE, Santa Cruz Biotechnology, sc-200800), BMS-345541 (Sigma, B9935) or DMSO. Cell extracts were subjected to Western blot analysis, as described previously. Quantification of eGFP protein from Western blot analysis was performed by densitometric analysis, using the Fiji (ImageJ) software, and was then normalised to the YopM-siCTL or empty vector-siCTL conditions.

**Immunofluorescence analyses**
Cells were fixed with 1X PBS-4% paraformaldehyde for 15 minutes at room temperature, followed by quenching (1X PBS-50 mM $NH_4Cl$) for 10 minutes. Cells were permeabilized with 1X PBS-0.1% Triton X-100 and blocked for 30 minutes with 1X PBS-5% BSA. Cells were then stained with DAPI (4',6-diamidino-2-phénylindole, dichlorhydrate; Invitrogen D1306) and Alexa Fluor™ 350 Phalloidin (Invitrogen; A22281) for 30 minutes at room temperature, followed by mounting to glass slides using Mowiol (Sigma). Immunosignals were analyzed with a Leica SP8 confocal microscope at 63X magnification. Images were processed using Fiji software. For colocalization analyses, individual cells were manually chosen as "regions of interest" (ROIs) within the deconvolved image through hand-drawing, the entire Z-stack was analyzed and the Manders' colocalization coefficient was obtained by using the Fiji JACoP Plugin (v2.1.4) (9).

**Western blot analyses**
Cells were washed with 1X PBS and lysed either in 1X Laemmli Loading Buffer or in Radio-ImmunoPrecipitation Assay (RIPA) buffer, for which the protein concentration was determined using the Bradford reagent (Bio-Rad, 5000006EDU). Approximately 100 μg of proteins were denatured 5 min at 95°C and then subjected to SDS-PAGE and Western blot analyses, according to standard procedures. The detection of proteins of interest was performed using the following primary antibodies: anti-Ago1 (D84G10, Cell Signaling Technology, 5053), anti-Ago2 (C34C6, Cell Signaling Technology, 2897), anti-Ago3 (4B1-F6, Active Motif, 39787), anti-Ago4 (D10F10, Cell Signaling Technology, 6913), anti-DDX6 (Bethyl, A300-460A), anti-Dicer (Cell Signaling Technology, 3363), anti-Flag (M2, Sigma-Aldrich, F1804), anti-GAPDH (14C10, Cell Signaling Technology, 2118), anti-GFP (D5.1, Cell Signaling Technology, 2956), anti-PABPC1 (Atlas Antibodies, HPA045423), anti-phospho-IκBα (Ser32) (14D4, Cell Signaling Technology, 2859), anti-Renilla (Abcam, ab185926), anti-TNRC6A (Bethyl, A302-329A) and anti-α-Tubulin (DM1A, Cell Signaling Technology, 3873). All antibodies were diluted 1/1000, excepted for anti-Renilla at 1/20 000. As secondary antibodies, anti-mouse (Cell Signaling Technology, 7076S) or anti-rabbit (Cell Signaling Technology, 7074S) were used and proteins were detected with Agrisera ECL SuperBright (Agrisera, AS16 ECL-S-100) using a LAS 4000 mini (GE Healthcare).

**Co-immunoprecipitation analyses**
Cells were transiently transfected in 10 $cm^2$ dishes, harvested at 48h post-transfection, washed twice with ice cold 1X PBS and lysed with Lysis Buffer (20 mM, Tris–HCl [pH 7.4], 150 mM NaCl, 2 mM EDTA, 0.5% NP-40, 1 mM DTT, 1X protease inhibitor [EDTA-free complete Protease Inhibitor Cocktail, Roche, 11873580001] and 1:100 phosphatase inhibitor [Sigma-Aldrich, P5726]). In some experiments, cell lysates were treated with 100 ng/μL of RNase A (ThermoFisher, R1253) for 1h at 37°C. The RNAse A treatment was checked on agarose gel by loading some input extract (data not shown). For each condition, 30 µL of input were collected and the concentration was measured using the Bradford reagent in order to load 100 µg of input proteins for the following analysis. Thirty µl of Dynabeads Protein G (Invitrogen, 10003D) were incubated with 7 μg of anti-FLAG M2 antibody (Sigma-Aldrich, F1804) for 1h30 at 4°C in 1X PBS containing 0.1% of Tween20. Dynabeads Protein G linked to the antibodies were then incubated overnight with cell lysates at 4°C under agitation. The immunoprecipitates were washed three times with IP Buffer (50 mM Tris–HCl (pH 7.4), from 150 mM to 250 mM NaCl, 0.05% NP-40, 1X protease inhibitor and phosphatase inhibitor) and eluted in 4X Laemmli Loading Buffer. Protein samples were denatured 5 min at 95°C, resolved on SDS-PAGE gel and analysed by Western blot.

**Purification of recombinant proteins**

All recombinant proteins were expressed from the *E. coli* strain BL21(DE3) codon plus (ThermoFisher, EC0114) in 1L of Terrific broth (TB). Cultures were grown at 37°C until they reached an $OD_{600}$ of 2 and protein production was induced with 1 mM Isopropyl-β-D-thiogalactopyranoside (IPTG), followed by overnight growth at 18°C with shaking at 180 rpm. Bacterial cells expressing $His_6$-SUMO-LegK1$^{2–386}$, $His_6$-SUMO-LegK1-KA$^{2–386}$, CBP-PIWI$^{517-859}$-$His_6$, CBP-IκBα-$His_6$ and $His_6$-TAP were collected by centrifugation, resuspended in Lysis Buffer (1.5X PBS, 1 mM $MgAc_2$, 0,1 % NP-40, 20 mM imidazole, 10 % glycerol), and lysed by sonication during 4 min on ice. Lysate was clarified by high-speed centrifugation (18,000 rpm) and then purified on 250 μL of Ni-NTA resin (Thermo Fisher Scientific, 88221). Resin was pre-equilibrated in Lysis Buffer, and supernatant was incubated with resin for 2h at 4°C. His fusion proteins linked to the resin were washed once with Lysis Buffer, then once with Wash Buffer (1.5X PBS, 250 mM NaCl, 1 mM $MgAc_2$, 0.1 % NP-40, 50 mM imidazole, 10 % glycerol), and finally with Lysis Buffer. Proteins were eluted in Elution Buffer (1.5X PBS, 1 mM $MgAc_2$, 0.1 % NP-40, 150 mM imidazole, 10 % glycerol). Excess imidazole was removed by overnight dialysis using Spectrum™ Labs Spectra/Por™ 2 12-14 kD MWCO Standard RC Dry Dialysis Kits (FisherScientific, 15310762) into Dialysis Buffer (1.5X PBS, 1 mM $MgAc_2$, 10 % glycerol, 2 mM DTT) before storage at -80°C.

Bacterial cells expressing GST-Ago2 recombinant protein were collected by centrifugation, resuspended in Lysis Buffer (1.5X PBS, 1 mM $MgAc_2$, 1mM DTT, 0.1% NP-40, 10% glycerol), and lysed by sonication during 4 min on ice. Lysate was clarified by high-speed centrifugation and then purified on 500 μL of Glutathione Sepharose 4B resin (Merck, GE-17-0756-01). Beads were pre-equilibrated in Lysis Buffer and supernatant was incubated with resin for 2h at 4°C. GST fusion proteins linked to the beads were washed once with Lysis Buffer, then once with Wash Buffer (1.5X PBS, 250 mM KCl, 1 mM $MgAc_2$, 1 mM DTT, 0.1 % NP-40, 10 % glycerol), and finally with Lysis Buffer. Proteins were eluted in Elution Buffer (50 mM Tris pH8.0, 250 mM KCl, 1 mM $MgAc_2$, 1 mM DTT, 0.1 % NP-40, 10 % glycerol, 10 mM reduced glutathione). Proteins were subjected to overnight dialysis using the same dialysis tubing as mentioned above into Dialysis Buffer (10 mM HEPES pH7.5, 250 mM KCl, 1 mM $MgAc_2$, 10 % glycerol, 2 mM DTT) before storage at -80°C. All recombinant protein concentrations were determined using the Bradford reagent. Proteins were analysed by Coomassie Blue staining after SDS-PAGE.

***In vitro* interaction assays**

Glutathione Sepharose 4B resin and Calmodulin Affinity Resin (Agilent Technologies, 214303) were washed twice with Blocking Buffer (20 mM Hepes pH 7.5, 150 mM NaCl, 0.1 % NP-40) and then blocked into Blocking Buffer supplemented with 20 mM NaCl, 2 mg/mL glycogen carrier, 10 mg/mL tRNA and 10 mg/mL BSA during 2h at 4°C. Subsequently, beads were washed twice with Blocking Buffer and resuspended into Storage Buffer (10 mM HEPES pH 7.5, 250 mM NaCl, 1 mM EDTA). For *in vitro* interaction assays using GST pull-down, 5 μg of each purified protein were incubated in B1.5 Buffer (1.5X PBS, 1 mM $MgAc_2$, 2 mM DTT, 10% glycerol). The same volume of 2X BB Buffer (40 mM Hepes pH 7.5, 83 mM NaCl, 1 mM $MgAc_2$, 0.2% NP-40, 11.7% glycerol, 2 mM DTT) was added. After 20 min of interaction at 30°C under rotation, 12 µL of Glutathione Sepharose 4B were added. Beads were washed three times with 500 µL of 1X BB 250/10 Buffer (20 mM HEPES pH 7.5, 250 mM NaCl, 1 mM $MgAc_2$, 0,2 % NP-40, 10 % glycerol, 1 mM DTT), and then proteins were eluted by incubation with 20 µL of Elution Buffer (50 mM Tris·HCl (pH 7.5), 250 mM KCl, 1 mM $MgAc_2$, 1 mM DTT, 0.1 % NP-40, 10 % glycerol, 10 mM reduced Glutathione) for 5 min at 30 °C, under shaking at 1,400 rpm. Finally, proteins bound to GST beads were eluted with Elution Buffer for 5 min at 30 °C, under shaking at 1,400 rpm. 4X Laemmli Loading Buffer was added and proteins were separated onto SDS-PAGE gel and analysed by Western blot.

For binding assays using CBP pull-down, proteins were prepared according to the same protocol as mentioned above. In some experiments, from 5 to 15 µg of human cell-extracted RNAs were incubated with protein samples. After 20 min of interaction at 30°C under rotation, 12 µL of Calmodulin Affinity Resin were added. Beads were washed three times with 500 µL of 1X BB 250/10 Buffer (20 mM Hepes pH=7.5, 250 mM NaCl, 2 mM $MgAc_2$, 2 mM imidazole, 2 mM $CaCl_2$, 0.1 % NP-40, 10 % glycerol, 1 mM DTT). Proteins were eluted by incubation with 20 µL of Elution

Buffer (10 mM Tris·HCl (pH 7.5), 250 mM KCl, 1 mM $MgAc_2$, 1 mM DTT, 0.1 % NP-40, 10 % glycerol, 20 mM EGTA) for 5 min at 30 °C, under shaking at 1,400 rpm. 4X Laemmli Loading Buffer was added and proteins were separated onto SDS/PAGE gel and analysed by Western blot.

**Kinase assays**

*In vitro* phosphorylation of 1 μg of purified GST-Ago2 or CBP-IκBα-$His_6$ recombinant proteins was performed in the presence of 1 μg of $His_6$-SUMO-LegK1$^{2–386}$ or $His_6$-SUMO-LegK1$^{2–386}$-KA in 20 μl of phosphorylation buffer, containing 25 mM Tris HCl (pH 7.5), 200 mM NaCl, 10 mM $MgCl_2$, and 50 μM unlabeled ATP. Reactions were split in two, and in one half 10 μCi [γ-32P] ATP were added. The phosphorylation reaction was performed for 1h at 30°C and stopped by adding 4X Laemmli Loading Buffer. A Western blot analysis was carried out on unlabeled samples. Labeled samples were analysed by SDS-PAGE, which was exposed to a phosphor screen and visualised by Typhoon FLA 9500 biomolecular imager.

The kinase activity of LegK1 mutant versions was determined using IκBα recombinant protein. For this, HEK293T cells were seeded in 6 $cm^2$ dishes and transfected with either pFlag-eGFP, p2xFlag-LegK1-WT, -KA, -3WF, -W283F, -W293F, -W293A or -W283F-W293A. Cells were lysed and subjected to a Flag immunoprecipitation. The resulting immunoprecipitates were incubated with Phosphorylation Buffer (250 mM Tris-HCl pH 7.5, 50 mM $MnCl_2$, 50 mM DTT) and 1 μg of purified CBP-IκBα-$His_6$ recombinant proteins for 30 min at 30°C. Proteins were mixed with 4X Laemmli Loading Buffer and then analysed by Western blot using the phospho-IκBα (Ser32) (14D4) antibody.

**Peptide pull-down assays**

Biotinylated peptides (synthetised by Genscript) were resuspended in DMSO, and heated at 40°C for 5 min. For binding assays, 10 µg of peptides were diluted into 200 µL of PBS containing 0.1% of Tween20, and were incubated for 30 min at room temperature in the presence of 30 µL of Dynabeads MyOne™ Streptavidin T1 (Thermo Fisher Scientific, 65601). The beads were then washed once in 0.1% PBS Tween and twice in IP Buffer (50 mM Tris-HCl pH 7.4, 150 mM NaCl, 5 mM $MgCl_2$, 0.05% NP-40, 1/100 Phosphatase inhibitor and 1X protease inhibitor). HEK293T cells were washed twice with ice cold 1X PBS and scraped with Lysis Buffer (see the Co-immunoprecipitation analyses section). Some cell lysate were collected and the concentration was measured using the Bradford reagent in order to load 80 µg of input proteins for the following analysis. Cells were incubated at 4°C under rotation for 30 min and the supernatant was harvested by centrifugation at 16,000 g for 10 min at 4°C. Beads-coupled peptides were incubated in the presence of cell lysates for 1 hour at 4°C After three washing steps of the beads with IP Buffer, the proteins were eluted in 4X Laemmli Loading Buffer and resolved on SDS-PAGE. The presence of Ago proteins was detected by Western blot analyses.

**RNA extractions and RT-qPCR**

For gene expression analyses, total RNAs were isolated by phenol-chloroform extraction using TRIzol Reagent (Thermo Fisher Scientific, 15-596-018), according to the manufacturer's instructions. Approximately 0.5 µg of RNAs were then digested by DNAse I (Promega, M6101) at 37°C for 50 min to remove the genomic DNA, followed by 10 min at 65°C to inactivate the DNAse. DNAse-digested RNAs were reverse-transcribed into complementary DNA using qScript cDNA Supermix (Quanta Biosciences, 733-1178). The cDNAs were quantified with SYBR Green qPCR mix (Takyon, Eurogentec, UF-NSMT-B0701) and gene-specific primers (*SI Appendix*, Table S1) in 384-well plates, following a protocol of heating at 95°C for 10 min, 45 cycles of denaturation at 95°C for 10 sec, and annealing at 60°C for 40 sec. A melting curve was performed at the end of the amplification, and transcript levels were then normalised to the abundance of *GAPDH* transcripts.

**Northern blot analyses**

Accumulation of low molecular weight RNAs was assessed by Northern blot analysis as previously described (10). RNA gel blot analysis of low molecular weight RNAs was performed on 30 µg of total RNAs. U6, let-7, miR16 and miR20a probes were generated using T4 Polynucleotide Kinase

(PNK; Thermo Scientific, EK0032) following the manufacturer instructions. DNA oligonucleotides complementary to miRNA sequences were end-labeled with γ-$^{32}$P-ATP using T4 PNK. The indicated probes were hybridized to the same membrane by sequential rounds of probing and stripping. Detection of U6 RNA was used to confirm equal loading.

**PRM data acquisition method**

HeLa cells were seeded in 15 cm$^2$ dishes at a density of 1.4x10$^7$ cells, and transiently transfected with either pFlag-eGFP, p2xFlag-LegK1-WT or -KA. At 24h post-transfection, cells were washed twice with ice cold 1X PBS and lysed with Lysis Buffer (see the Co-immunoprecipitation analyses section). For each condition, 30 µL of input were collected and analysed by Western blot. Sixty µl of Dynabeads Protein G were incubated for 1h30 at 4°C under agitation with 16 µg of the anti-Ago2 antibody (Sigma-Aldrich, SAB4200085) in 1X PBS containing 0.1% of Tween 20. Dynabeads Protein G linked to Ago2 antibody were then incubated overnight with cell lysates at 4°C under agitation. The immunoprecipitates were washed three times with IP Buffer (see the Co-immunoprecipitation analyses section) and eluted in 4X Laemmli Loading Buffer. Five immunoprecipitations of each condition were performed. Protein samples were denatured 5 min at 95°C, then were simultaneously separated on SDS-PAGE and stained with colloidal blue (LabSafe Gel Blue, Gbiosciences, 786-35). One gel slice was excised for each purification and in-gel digested by using trypsin/LysC (Promega, V5072). Peptides extracted from each band were then loaded onto a homemade C18 StageTips for desalting. Peptides were eluted using 40/60 MeCN/$H_2O$ + 0.1 % formic acid and vacuum concentrated to dryness. Peptide samples were resuspended in Buffer A (2/98 MeCN/$H_2O$ in 0.1 % formic acid), separated and analysed by nanoLC-MS/MS using an RSLCnano system (Thermo Scientific, Ultimate 3000) coupled online to a Q Exactive HF-X mass spectrometer (Thermo Scientific). Peptides were first trapped onto a C18 column (75 µm inner diameter × 2 cm; nanoViper Acclaim PepMapTM 100, Thermo Scientific) with Buffer A at a flow rate of 2.5 µL/min over 4 min. Separation was performed on a 50 cm x 75 µm C18 column (Thermo Scientific, nanoViper C18, 3 µm, 100Å, Acclaim PepMapTM RSLC) regulated to 50°C and with a linear gradient from 2% to 30% of buffer B (100% MeCN in 0.1% formic acid) at a flow rate of 300 nL/min over 91 min. The mass spectrometer was operated in Parallel Reaction Monitoring (PRM) mode (see acquisition list Supplementary Table 2). The acquisition list was generated from the peptides obtained from the mix samples (5 replicates) of each condition, based on the data-dependent acquisition (DDA) results (data not shown).

**PRM data analyses**

The PRM data were analysed using Skyline 4.1 (MacCoss Lab Software, Seattle, Washington [https://skyline.ms/project/home/software/Skyline/begin.view]). Fragment ions for each targeted mass (*SI Appendix*, Table S3) were extracted and peak areas were integrated. The peptide areas were log2 transformed and the mean log2- area was normalised by the mean area of five Ago2 peptides (ELLIQFYK, SGNIPAGTTVDTK, SIEEQQKPLTDSQR, VELEVTLPGEGK and VLQPPSILYGGR) using software R 3.1.0. On each phospho-peptide, a linear model was used to estimate the mean fold change between the conditions, its 95% confidence interval and the p-value of the two-sided associated t-test. The p-values were adjusted with the Benjamini-Hochberg procedure (11). Data are available *via* ProteomeXchange with identifier PXD037279 (Username:; Password: xzomxsb6) (12).

**Prediction of W-motifs**

W-motifs were predicted using the Wsearch algorithm (13), which scores the W-motifs according to a Position-Specific Scoring Matrix (PSSM) derived from experimentally validated Ago-binding motifs from animals. Briefly, the protein sequences of all bacterial organisms tested were retrieved from Uniprot (14) and subjected to Wsearch either on the website or using the python script. The score of the different positive motifs were then added together, corresponding to the final score. For the first pre-selection, we applied an arbitrary cut-off greater than six on the average W-scores of all motifs present in the candidate protein sequence. Among the bacterial candidate proteins exhibiting the highest W-score, we further selected known secreted virulence factors or putative virulence factors predicted to be secreted using PSORTb 3.0 (https://www.psort.org/psortb/) (15).

**Predicting the effect of W-motif mutations on LegK1 protein stability**

The coordinates of LegK1 models were generated using AlphaFold (version 2.3.1) via ColabFold (version 1.5.2) (16, 17). The five structural models were superimposed and analyzed using PyMOL (https://www.pymol.org/). In order to predict the effect of W-motif mutations on the stability of LegK1, PoPMuSiC (v3.1) and HoTMuSiC (v1.0) softwares were used (18, 19). The coordinates of LegK1 models were provided to both PoPMuSiC and HoTMuSiC software as input. For each mutation, the average value as well as the standard deviation for the set of predicted values obtained for each of the five LegK1 models are reported. Mutations tested correspond to W41F, W283F, W293F, W293A.

**Alignment of orthologous protein sequences and phylogeny**

The protein sequence of LegK1 (from the *lpp1439* gene) from *L. pneumophila* strain Paris was used as a reference to retrieve homologous LegK1 protein sequences from the order *Legionellales* (taxid: 445). To this end, a BLASTP (Basic Local Alignment Search Tool) was performed on the NCBI (National Center for Biotechnology Information) website. An identity cut-off of 40%, an Expect (E)-value cut-off of $10^{-5}$ and a minimum percentage match length of subject and query of 65% were used. The set of homologous protein sequences were then aligned using ClustalW2 on Geneious 10.2.6 software. Tree was conducted using Neighbor-joining.

**Heatmap of *Ago* transcripts**

The RNA-sequencing results were generated in the Cancer Cell Line Encyclopedia (CCLE, Primary ID: E-MTAB-2770) and are reported as linear values, retrieved from Genevestigator.

**Quantification and statistical analysis**

Experimental results were shown as standard error of the mean (SEM) for luciferase reporter and qPCR experiments and as standard deviation (SD) for infection assays. Statistical analysis was carried out using the Prism Software (GraphPad Prism 9.0). For statistical comparison between two groups or several conditions, an unpaired t test or one-way analysis of variance (ANOVA) of biological replicates were used, respectively. A value of $P < 0.05$ was considered statistically significant. All statistical tests are specified in the respective figure legends.

### Supporting Information Text

#### LegK1 does not phosphorylate human Ago2

Since LegK1 can physically interact with human Ago2 and possesses a eukaryotic-like serine/threonine kinase activity, we reasoned that it might directly phosphorylate Ago2 residues to inhibit its functions. To test this hypothesis, we first analyzed specific phosphorylation sites on Ago2 through a mass spectrometry targeted approach. More specifically, we transfected the pFlag-eGFP, p2xFlag-LegK1-WT or p2xFlag-LegK1-KA plasmids in HEK293T cells and further subjected the corresponding cell lysates to Ago2 immunoprecipitation and Parallel Reaction Monitoring (PRM) data acquisition method (*SI Appendix*, Fig. S7*B-E*, Table 2 and 3). The latter approach relies on a liquid chromatography-mass spectrometry (LC-MS)-based targeted peptide quantification method. In more details, Ago2 immunoprecipitates were separated by SDS-PAGE, gel slices were excised for each purification, and subsequently in-gel digested by trypsin. Digested peptides were then analyzed by LC-MS/MS on a Q Exactive HF-X mass spectrometer. Results from this analysis allowed us to retrieve four phosphopeptides and to quantify the phosphates on the serine residues at positions 387, 824 and 828 (*pS*) of Ago2. However, we did not observe a significative differential phosphorylation status (ratio > 2) of these targeted serine residues in the presence of LegK1 or LegK1-KA compared to the control eGFP condition (*SI Appendix*, Fig. S7*F*, Table 2 and 3). These data indicate that LegK1 does not interfere with the phosphorylation status of those Ago2 serine residues. However, this approach was limited to the phosphorylation of the serine residues from the recovered phosphopeptides in the samples and did not provide information on the phosphorylation status of all the remaining serine and threonine residues of Ago2. As a complementary approach, we further decided to perform an *in vitro* kinase assay in the presence of the truncated LegK1$^{2:386}$ or LegK1$^{2:386}$-KA versions. Because I$\kappa$B$\alpha$ has previously been shown to be phosphorylated by LegK1 on the serine residues 32 and 36, we used this host target as an internal control for LegK1 phosphorylation. By incubating purified CBP-I$\kappa$B$\alpha$-His$_6$ with His$_6$-SUMO-LegK1$^{2:386}$ proteins, we found a clear incorporation of $^{32}$P radioactivity (*SI Appendix*, Fig. S7*G*), confirming previous findings. In contrast, we did not find any incorporation of $^{32}$P radioactivity upon incubation of purified GST-Ago2 with His$_6$-SUMO-LegK1$^{2:386}$ proteins, as observed upon incubation of purified His$_6$-TAP control proteins with His$_6$-SUMO-LegK1$^{2:386}$ (*SI Appendix*, Fig. S7*G*). Collectively, these data indicate that LegK1 does not phosphorylate human Ago2 *in vitro*.

### Supplementary Figures

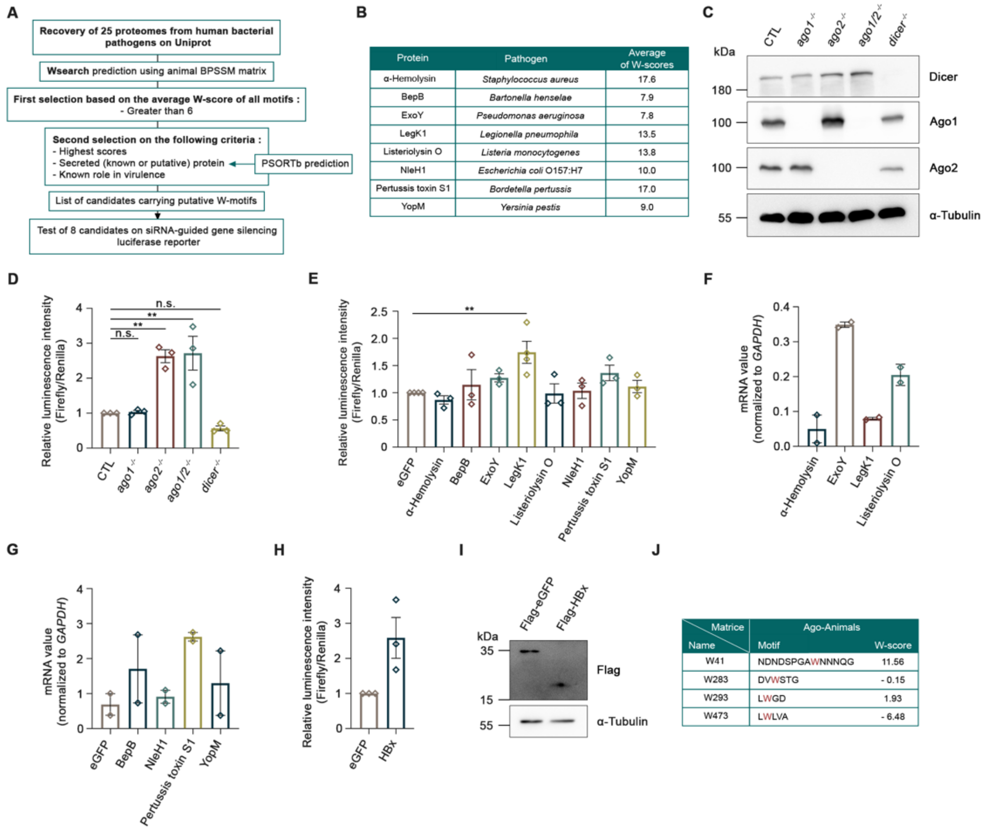


**Fig. S1. Screening of candidate bacterial proteins carrying W-motifs on the Ago2-dependent *CXCR4* siRNA silencing reporter**

(*A*) *In silico* approach used to identify bacterial proteins possessing canonical W-motifs. Wsearch algorithm is based on a Bidirectional Position-Specific Scoring Matrice (BPSSM). The source sequence dataset consists of experimentally validated W-rich protein domains. The animal matrix was generated from the orthologous sequences of TNRC6A, TNRC6B, TNRC6C (from mammals), GAWKY (from fruit fly), AIN1, AIN2 (from worm) and PrP (from prion). PSORTb is a bacterial localization prediction program that was used to identify predicted secreted proteins. (*B*) Scores of predicted W-motifs on a selection of bacterial candidate proteins. The sum of Wsearch scores of the predicted W-motifs from each bacterial protein was determined using the Ago-Animals matrix of the Wsearch algorithm, as described in (*A*). (*C*) Immunoblot of Ago and Dicer proteins in control (CTL), *ago1*$^{-/-}$, *ago2*$^{-/-}$, *ago1*$^{-/-}$*ago2*$^{-/-}$, and *dicer*$^{-/-}$ HeLa cell lines. α-Tubulin was used as a loading control. (*D*) The CXCR4 siRNA silencing reporter is dependent on Ago2 proteins. Control (CTL), *ago1*$^{-/-}$, *ago2*$^{-/-}$, *ago1*$^{-/-}$*ago2*$^{-/-}$, and *dicer*$^{-/-}$ HeLa cell lines were co-transfected with CXCR4 luciferase-based siRNA silencing reporter and control siRNA (siCTL) or *CXCR4* siRNA duplexes. The luminescence intensity of Firefly luciferase relative to the one of Renilla was calculated, normalized to the siCTL condition, and further normalized to the CTL HeLa cell condition. (*E*) The LegK1 effector suppresses CXCR4 siRNA silencing reporter. HeLa cells were co-transfected with CXCR4 luciferase-based siRNA silencing reporter, control siRNA (siCTL) or *CXCR4* siRNA duplexes and vectors expressing Flag-eGFP or the bacterial candidates. Luciferase expression was measured at 48h post-transfection. The luminescence intensity of Firefly luciferase relative to the one of Renilla was calculated, normalized to the siCTL condition, and further normalized to the eGFP condition. *(F)* and *(G)* mRNA values of (*F*) low expressed or (*G*) normally expressed candidate bacterial genes in human cells. HeLa cells were transfected with a plasmid expressing each candidate bacterial gene or the control eGFP for 48 hours. The RT-qPCR data are represented as absolute values that were normalized to the absolute value of the constitutively expressed *GAPDH* gene. It is noteworthy that, although the candidate bacterial effectors were all expressed at the mRNA levels, only listeriolysin O, NleH1 and pertussis toxin S1 were readily detected at the protein levels. This phenomenon is commonly observed with bacterial effectors which cognate sequences have not been subjected to codon optimization. Therefore, we do not exclude the possibility that the absence of siRNA reporter derepression for these candidate effectors, could simply be due to their absence or low abundance in the cells. (*H*) Luciferase-based siRNA silencing reporter assay in the presence of the VSR HBx in HeLa cells. The experiment was performed as described in (*E*), but the vectors expressing Flag-eGFP or 3xFlag-HBx were co-transfected along with the reporter system. The relative Firefly/Renilla value was normalized to that of the eGFP condition. (*I*) Immunoblot of control eGFP and HBx proteins in HeLa cells. Cells were transiently transfected with vectors expressing either Flag-eGFP or 3xFlag-HBx constructs for 48 hours. α-Tubulin was used as a loading control. (*J*) Scores of the predicted W-motifs present in the LegK1 protein sequence. The Wsearch scores were determined using the Ago-Animals matrix of the Wsearch algorithm, as described in (*A*). Data are represented as ± SEM. Data are representative of two (*F* and *G*) or three (*D*, *E* and *H*) independent experiments.

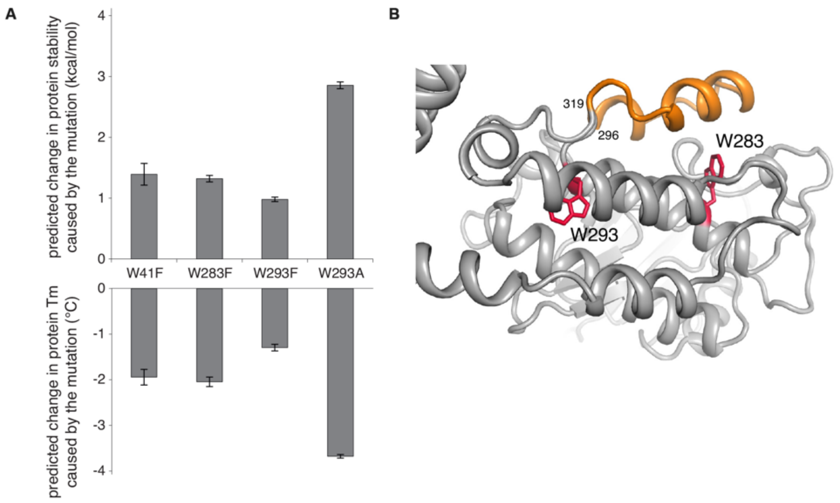


**Fig. S2. Prediction of changes in protein stability caused by punctual mutations of W-motifs and the position of W-motifs**

(*A*) Prediction of changes in protein stability caused by protein mutations as reported by PoPMuSiC (v3.1) and HoTMuSiC (v1.0) softwares. Predicted changes in Gibbs free energy of folding (ΔΔG) in kcal/mol are reported in the upper part of the graph, and predicted changes in protein melting temperature (ΔTm) in °C are reported in the lower part. Positive ΔΔG and negative ΔTm correspond to destabilizing mutations. For each mutation (*i.e*. W41F, W283F, W293F, W293A), the average value as well as the standard deviation for the set of predicted values for identical mutations in the ensemble composed of five ColabFold models of LegK1 are reported. (*B*) Cartoon representation of the best ranked ColabFold model of LegK1 with residues W283 and W293 of W-motifs shown as red sticks. A small protein segment, *i.e*. residues 296-319, covering the W-motif W283, is shown in orange.

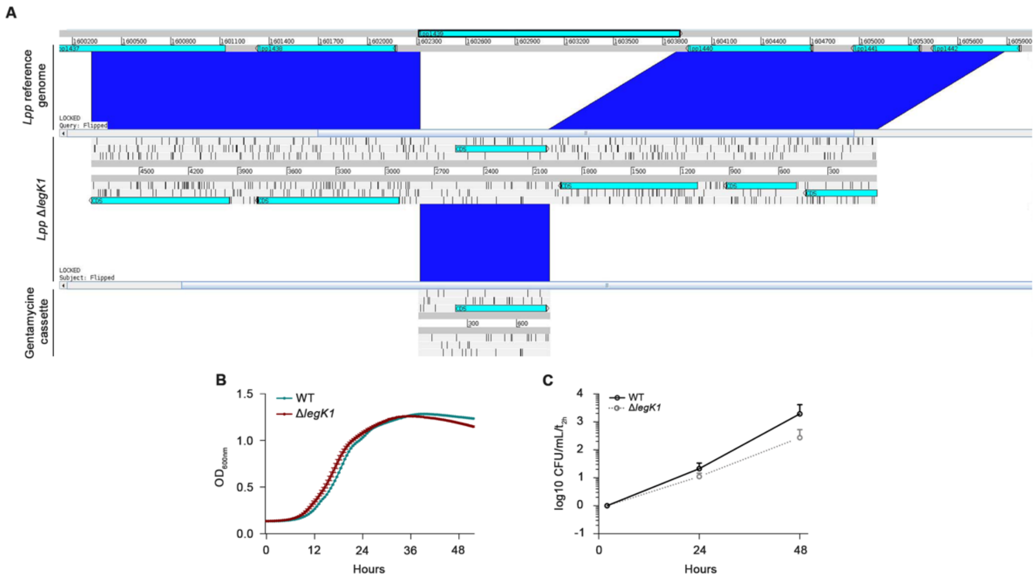


**Fig. S3. Characterization of the *Lpp* Δ*legK1* strain *in vitro* and in HK293-FcγRII**
(*A*) Screenshot depicting the specific deletion of the *legK1* gene and replacement with a gentamycin resistance gene in the *Lpp* Δ*legK1* strain. (*B*) Growth of *L. pneumophila* WT and Δ*legK1* strains was determined by measuring $OD_{600nm}$ over 48 hours at 37°C under shaking. (*C*) Intracellular replication of the *Lpp* WT or Δ*legK1* strains in HEK293-FcɣRII cell line. Cells were infected with *Lpp* WT strain at a MOI of 20 and bacterial titers were monitored at 24 and 48 hours post-infection (*hpi*). Results are shown as log10 ratio CFU/mL, where CFUs were normalized with the associated condition at 2h post-infection, corresponding to the entry of bacteria in host cells. Data are representative of three (*B* and *C*) independent experiments.

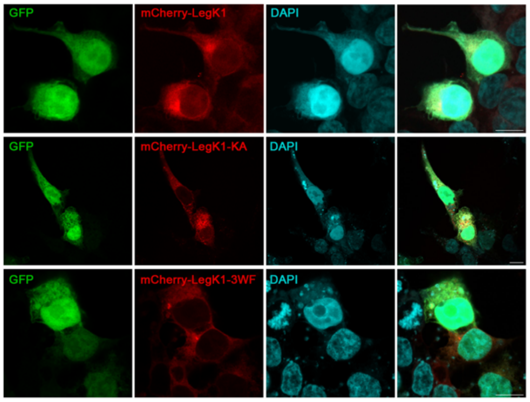


**Fig S4. Microscopy images showing the absence of colocalization of GFP control with WT and LegK1 mutants**

Representative confocal microscopy images of HEK293-FcγRII cells transfected for 48 hours with eGFP together with mCherry-LegK1 and its mutated forms, as indicated. DAPI are in cyan. Single channel and merged images are shown. Z stack images were acquired and deconvolution images are presented as maximum intensity z-projection. Scale bars 10 µm.

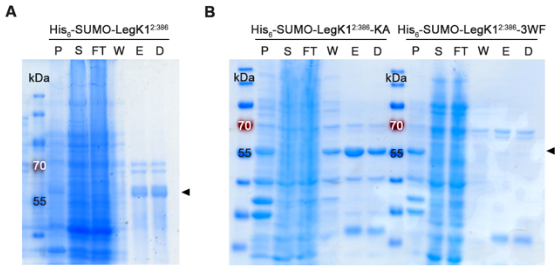


**Fig S5. Coomassie Blue stained analytical SDS-PAGE showing the fractions of the LegK1 WT, LegK1-KA and LegK1-3WF recombinant proteins purification in *E. coli***

(*A*) His$_6$-SUMO-LegK1 (a.a. 2:386), (*B*) His$_6$-SUMO-LegK1$^{2:386}$-KA and His$_6$-SUMO-LegK1$^{2:386}$-3WF recombinant proteins were expressed in *E. coli* strain BL21 (DE3) codon plus and purified by affinity chromatography. Expressions were induced with IPTG, followed by overnight growth, and bacterial cells were collected by centrifugation, resuspended and lysed by sonication. Lysates were centrifuged in a high-speed centrifuge to separate the pellets (P) from the supernatants (S). The supernatants were incubated with resin for 2h, subsequently the flow-throughs (FT) were removed. The protein-bound resin was washed (W) and proteins were eluted from the beads (E). The imidazole excess was removed by overnight dialysis (D). Arrows indicate His$_6$-SUMO-LegK1 WT and kinase-dead mutant recombinant proteins.

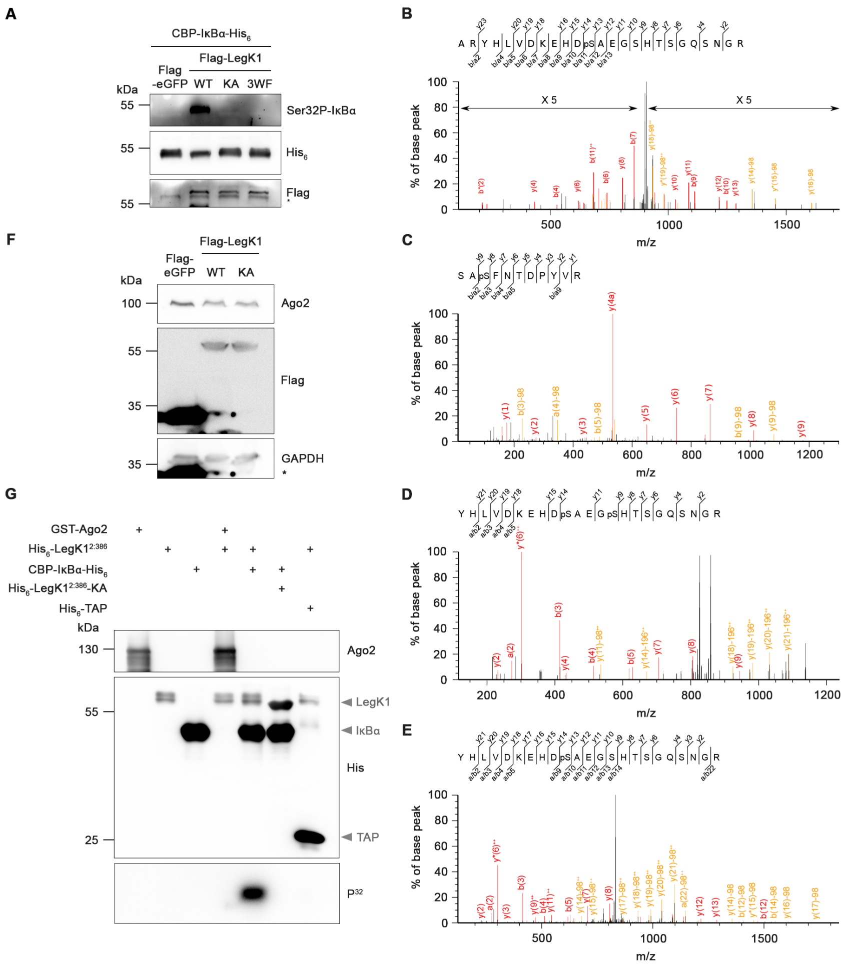


**Fig. S6. The Ago-binding platform of LegK1 is required for its kinase activity, but this effector does not phosphorylate human Ago2**

(*A*) The Ago-binding mutant of LegK1 does not phosphorylate the serine 32 of IκBα. Kinase assay in the presence of LegK1 WT, LegK1-KA or LegK1-3WF mutants. HEK293T cells were co-transfected with expression vectors for eGFP, 2xFlag-LegK1, 2xFlag-LegK1-KA or 2xFlag-LegK1-3WF. At 48h post-transfection, cells were lysed and subjected to a Flag immunoprecipitation. The resulting immunoprecipitates were incubated with purified CBP-IκBα-$His_6$ recombinant protein. The phosphorylation of the serine 32 of IκBα was analysed by Western blot using a specific antibody. * represents a specific band. The results shown are representative of two independent experiments. *(B-E)* Quantification of Ago2 phosphorylation by Parallel-Reaction Monitoring (PRM) approach using Liquid Chromatography with tandem mass spectrometry (LC-MS/MS) on selected serine residues. HEK293T cells were transfected with vectors expressing Flag-eGFP, 2xFlag-LegK1 or 2xFlag-LegK1-KA. At 24h post-transfection, endogenous Ago2 proteins were immunoprecipitated using an anti-Ago2 antibody. Representative HCD fragmentation spectra of human Ago2 phosphorylated-peptides obtained on a Q Exactive HF-X mass spectrometer are shown. The spectrum represents the peptide sequence and the observed ions of the phospho-peptide. (*B*) MS/MS of the tryptic (2 missed cleavages) ARYHLVDKEHD*pS*AEGSHTSGQSNGR (m/z 940.083+) peptide with the position of the phosphate group at S284 (mascot score 75.2) and labeled to show singly and doubly charged b and y ions, as well as ion corresponding to neutral losses of NH3 (*) and H3PO4 group (98Da). (*C*) MS/MS of the tryptic SA*pS*FNTDPYVR (m/z 668.782+) peptide with the position of the phosphate group at S387 (mascot score 57.2) and labeled to show singly charged a, b and y ions, as well as ion corresponding to neutral losses of H3PO4 group (98Da). (*D*) MS/MS of the tryptic (1 missed cleavage) YHLVDKEHD*pS*AEGpSHTSGQSNGR (m/z 891.023+) peptide with the position of the phosphate group at S824 and S828 (mascot score 24.9) and labeled to show singly and doubly charged a, b and y ions, as well as ion corresponding to neutral losses of NH3 (*) and H3PO4 group (98Da). (*E*) MS/MS of the tryptic (1 missed cleavage) YHLVDKEHD*pS*AEGpSHTSGQSNGR (m/z 864.373+) peptide with the position of the phosphate group at S824 (mascot score 82.5) and labeled to show singly and doubly charged a, b and y ions, as well as ion corresponding to neutral losses of NH3 (*) and H3PO4 group (98Da). (*F*) Immunoblot of cell lysates used for mass spectrometry analysis. HeLa cells were transfected with vectors expressing Flag-eGFP, 2xFlag-LegK1 or 2xFlag-LegK1-KA. At 24h post-transfection, total cells were lysed and were analysed by Western blot using indicated antibodies. The data shown is representative of five independent experiments. GAPDH was used as a loading control. * represents a specific band. *(G) In vitro* kinase assay of LegK1 in the presence of Ago2. Purified $His_6$-SUMO-$LegK1^{2:386}$ or $His_6$-SUMO-LegK1-$KA^{2:386}$ recombinant proteins were incubated with [γ-$P^{32}$] ATP in presence of GST-Ago2, CBP-IκBα-$His_6$ or $His_6$-TAP, as negative control. Incorporation of phosphate into the protein was examined by autoradiography of the reaction mixtures resolved on the SDS-PAGE gel, as shown. Loading amounts of the different recombinant proteins were assessed by Western blot analysis using indicated antibodies. Blots are representative of three independent experiments.

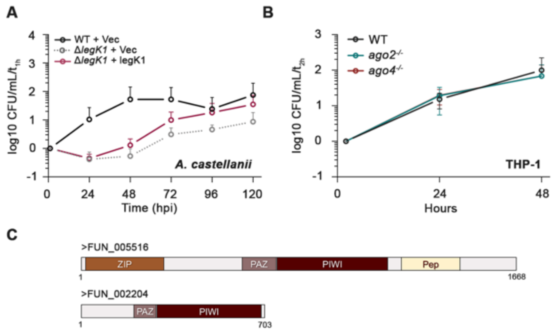


**Fig. S7. Characterization of the *Lpp* Δ*legK1* strain in *A. castellanii* and in *ago2*$^{-/-}$ or *ago4*$^{-/-}$ THP-1 cells**

(*A*) Complementation assays in *A. castellanii.* Cells were infected at a MOI of 1 at 20°C with *Lpp* WT or Δ*legK1* mutant strains carrying the empty vector pBC-KS (Vec) or the complementing plasmid pBC-KS-LegK1*.* Intracellular replication was determined by recording the number of colony-forming units (CFU) by plating on buffered charcoal yeast extract (BCYE) agar. Results are shown as log10 ratio CFU/mL, where CFUs were normalized with the associated condition at 1h post-infection, corresponding to the entry of bacteria in host cells. (*B*) Intracellular replication of the *Lpp* WT strain in THP-1 monocyte-derived macrophages depleted of *Ago2* or *Ago4*. WT*, ago2* $^{-/-}$ and *ago4* $^{-/-}$ THP-1 macrophages were infected with the *Lpp* WT strain at a MOI of 10 and bacterial titers were monitored at 24- and 48-hours post-infection (*hpi*). Results are shown as log10 ratio CFU/mL, where CFUs were normalized with the associated condition at 2h post-infection, corresponding to the entry of bacteria in host cells. Data are represented as ± SD. Data are representative of three independent experiments. Two replicates from plotted data of infections with *Lpp* WT are the same data shown in Fig. 6*B*. (*C*) Schematic representation of Ago-like proteins identified in *A. castellanii C3* strain. PAZ, Piwi Argonaute and Zwille; Pep, peptidase C19; ZIP, Zrt/Irt-like proteins.

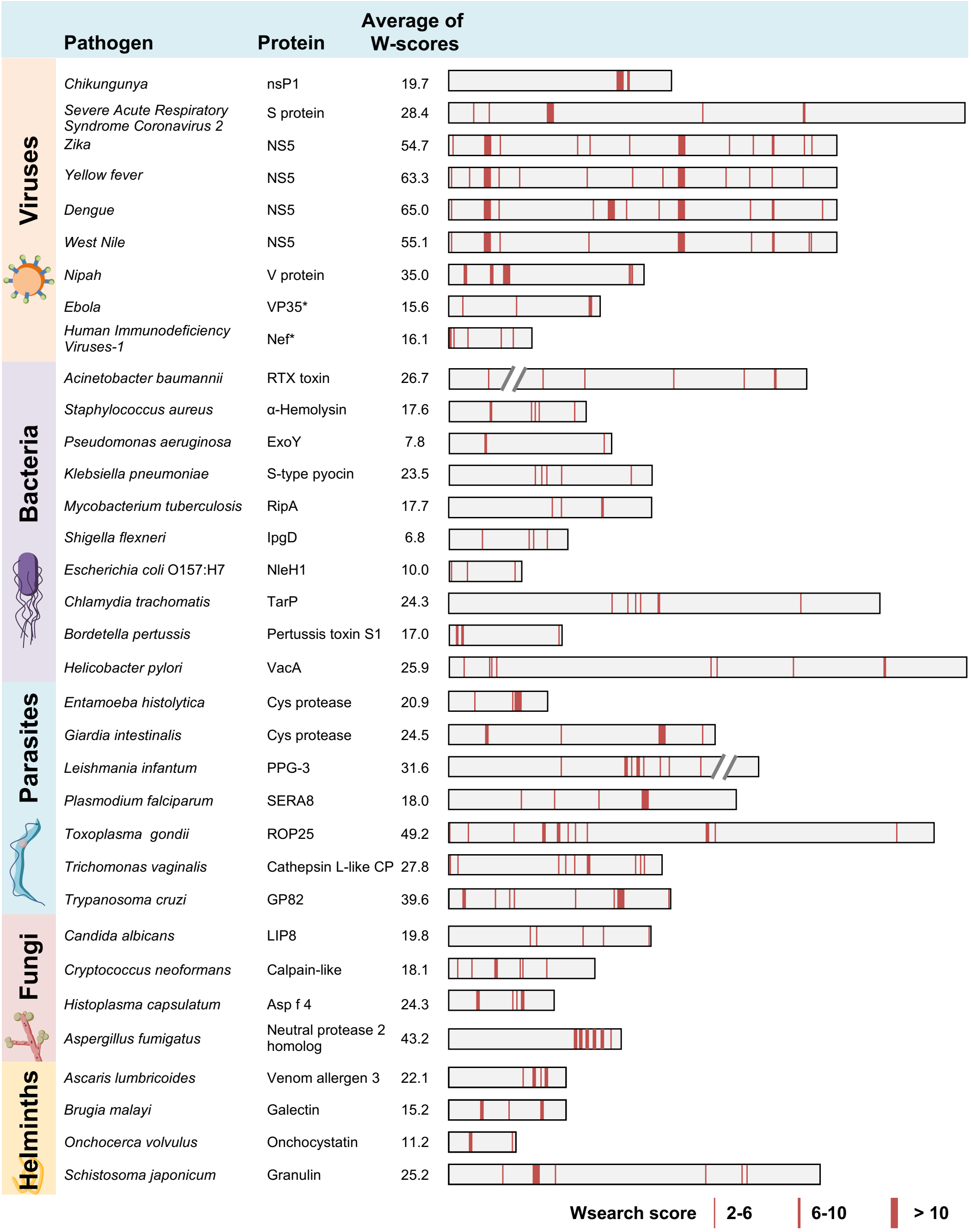


**Fig. S8. Scores of predicted W-motifs on a selection of virulent factors from viruses, non-viral pathogens and parasites**

The Wsearch scores sum of the predicted W-motifs from each human pathogenic non-viral and viral protein was determined using the animal (Ago-Animals) and viral (Ago-Plantvir) matrices of the Wsearch algorithm, respectively. Wsearch uses Position-specific Scoring Matrices (PSSM) to score W-containing sequences. The source sequence dataset consists of experimentally validated W-rich protein domains. The animal matrix was generated from the orthologous sequences of TNRC6A, TNRC6B, TNRC6C (from mammals), GAWKY (from fruit fly), AIN1, AIN2 (from worm) and PrP (from prion). The viral matrix was generated from the P1, NSs, P38, P37 proteins from 15 plant viruses. * corresponds to VSRs that have been previously experimentally validated.

**Supplementary Tables**

**Table S1. Primer sequences**

| **Plasmid construction** | | |
|---|---|---|
| **Oligo Name** | **Sequence (5' to 3')** | **Note** |
| BepB-GW_F | CACCATGCCAAAAGCAAAAGCAA | |
| BepB-stp-R | TTAGCTGGCAATAGCAAGCG | |
| Cassette_F | GATGAAGGCACGAACCCAGTTGACA | *Lpp* Δ*legK1* mutant |
| Cassette_R | CGGCTTGAACGAATTGTTAGGTGGC | *Lpp* Δ*legK1* mutant |
| ExoY-GW_F | CACCATGCGTATCGACGGTCAT | |
| ExoY-stp-Rev | TCAGACCTTACGTTGGAA | |
| FinalPCR_F | CGGTCATGCTAAATGATTCAAGAATG | *Lpp* Δ*legK1* mutant |
| FinalPCR_R | CTTCATTCACTACCCAATTTAATACTGAATTGG | *Lpp* Δ*legK1* mutant |
| Flag2-pDEST-F2 | CATGATATTAAGTACCCTTATGACGTGCCCGATTAC | Introduction of one Flag |
| FLAG2-pdest-R | AGGGTACTTAATATCATGATCCTTGTAGT | Introduction of one Flag |
| HLA_F | CACCATGGCAGATTCTGATATTAACATTAAAACC | |
| HLA_R | TTAATTTGTCATTTCTTCTTTTTCCCA | |
| IκBα-BamHI_F | gagcGGATCCATGTTCCAGGCGGCCGAG | |
| IκBα-HindIII_R | ggtaAAGCTTCTCGTCCTCTGTGAACTCC | |
| LegK1-downstream_F | GCCACCTAACAATTCGTTCAAGCCGGTATTGCGGTTGGAAATTAAAAATCT | *Lpp* Δ*legK1* mutant |
| LegK1-downstream_R | CTTCATTCACTACCCAATTTAATACTGAATTGG | *Lpp* Δ*legK1* mutant |
| LegK1-K121A-F | TTAGTTGCCATACAAAACCATAGTGAACGC | |
| LegK1-K121A-R | GTTTTGTATGGCAACTAATTTATTCTTTGGTGG | |
| LegK1-upstream_F | CGGTCATGCTAAATGATTCAAGAATG | *Lpp* Δ*legK1* mutant |
| LegK1-upstream_R | TGTCAACTGGGTTCGTGCCTTCATCCGAGGCATGATATTGACTTCAATTT | *Lpp* Δ*legK1* mutant |
| LegK1-W283F_F | GATGTTTTCTCAACAGGCAGGATATTAAG | |
| LegK1-W283F_R | TGTTGAGAAAACATCAGCTTTTGCTGTGT | |
| LegK1-W293A_F | TATTTGGCCGGTGATAAATATACTAATTATT | |
| LegK1-W293A_R | ATCACCGGCCAAATAACTTAATATCCTGC | |
| LegK1-W293F_F | TATTTGTTCGGTGATAAATATACTAATTATT | |
| LegK1-W293F_R | ATCACCGAACAAATAACTTAATATCCTGC | |
| LegK1-W41F-F | GTGCCTTCAATAATAATCAAGGATACAAAT | |
| LegK1-W41F-R | ATTATTGAAGGCACCTGGTGAATCATTATC | |
| LegK1-XhoI_F | CTCGAGCTATGCCTCGTACAATGTTTTTTTC | pmCherry-C1 |
| LegK1-BamHI_R | GGATCCTTAATTTCCAACCGCAATACC | pmCherry-C1 |
| LLO_F | CACCATGAAAAAAATAATGCTAGTGTTTATTACAC | |
| LLO_R | TTATTCGATTGGATTATCTACTTTATTA | |
| Lpp1439-promoteur_F | GGGCGAGCTCGAGTGTTTTCATCTCTTTCAATTG | pBC-KS-LegK1 |
| Lpp1439-promoteur_R | GGCTGGTACCTTAATTTCCAACCGCAATACCCA | pBC-KS-LegK1 |
| NleH1-GWF | CACCATGCTATCACCATCTTCT | |
| NleH1-stp-Rev | CTAAATTTTACTTAATACCACACT | |

| | | |
|---|---|---|
| PIWI-hAgo2-BamHI_F | GAGCGGATCCATGCTGGTGGTGGTCATCCTGC | |
| PIWI-hAgo2-HindIII_R | GGTAAAGCTTCAGGTGGTACCTGGCCC | |
| PT_F | CACCATGCGTTGCACTCGGGCAA | |
| PT_R | CCAGGTCTAGAACGAATACG | |
| T6B_F | CACCATGGATTGTCAGGCTGTCTTGCAGAC | |
| T6B-nostp_R | GAGCTCCCCCCATCCAGACT | |
| YopM-GWF | CACCATGTTCATAAATCCAAGAAAT | |
| YopM-stp-Rev | CTACTCAAATACATCATCTTC | |
| **qPCR** | | |
| **Oligo Name** | **Sequence (5' to 3')** | |
| BepB-RT_F | GCCACACTCCCTTCACATTT | |
| BepB-RT_R | TTTTTCGGCAAGCTCTTGAT | |
| eGFP-RT_F | GACGTAAACGGCCACAAGTT | |
| eGFP-RT_R | GAACTTCAGGGTCAGCTTGC | |
| ExoY-RT_F | CGAATACCGTCTTTGGCATT | |
| ExoY-RT_R | AGTTCGAGCTTTTCCCCTTC | |
| GAPDH-RT_F | GAACATCATCCCTGCCTCTACT | |
| GAPDH-RT_R | ATTTGGCAGGTTTTTCTAGACG | |
| HLA-RT_F | AACGAAAGGAACCATTGCTG | |
| HLA-RT_R | AAGGCCAGGCTAAACCACTT | |
| LegK1-RT_F2 | CACCGAGATGCCTTATGGTT | |
| LegK1-RT_R2 | GCTTTTCATTTCCTCCACCA | |
| LLO-RT_F | CGTCCATCTATTTGCCAGGT | |
| LLO-RT_R | ATTTCGGATAAAGCGTGGTG | |
| NleH1-RT_F | GATTATGCACCGCCAGAGTT | |
| NleH1-RT_R | CTTTTGTTGCATCCTCAGCA | |
| PT-RT_F | CAGAGCGAATATCTGGCACA | |
| PT-RT_R | AATACTCCGTGGTCGTGGTC | |
| YopM-RT_F | **AGCGTTACCTCCACGCTTAG** | |
| YopM-RT_R | GGAGCTGTTTCAGGTTTTGC | |
| **Probes for Northern blot** | | |
| **Oligo Name** | **Sequence (5' to 3')** | **Reference miRbase** |
| hsa-let-7a-5p | AACTATACAACCTACTACCTCA | MIMAT0000062 |
| hsa-miR-16-5p | CGCCAATATTTACGTGCTGCTA | MIMAT0000069 |
| hsa-miR-20a-5p | CTACCTGCACTATAAGCACTTTA | MIMAT0000075 |
| U6 | AGGGGCCATGCTAATCTTCTC | |

**Table S2. PRM targeted peptides using mass spectrometry**

| Peptide Sequence and Modifications position | Phosphorylated residue | Precursor Mass MH+ | Charge | Extracted fragments |
|---|---|---|---|---|
| ARYHLVDKEHDS[Phospho]AEGSHTSGQSNGR | S824 | 940,081146 | 3 | y14-98, y11, y9, y8, y7, y18(2+), y18-98(2+), y16-98(2+), y14-98(2+), b7, b9, b11, b11(2+), b14-98(2+) |
| SAS[Phospho]FNTDPYVR | S387 | 668,782099 | 2 | y9-98, y8, y7, y6, y5, y4, y9-98(2+), b2, b3-98, b4-98, b5-98, b7-98 |
| YHLVDKEHDS[Phospho]AEGS[Phospho]HTSGQSNGR | S824 & S828 | 891,023849 | 3 | y7, y6, y21-98(2+), y20-98(2+), y19-98(2+), y18-98(2+), y14-98(2+), y22-98(3+), b2, b3, b4, b5, b9, b21(3+) |
| YHLVDKEHDS[Phospho]AEGSHTSGQSNGR | S824 | 864,368405 | 3 | y11, y8, y7, y6, y21(2+), y21-98(2+), y20(2+), y20-98(2+), y19-98(2+), y18-98(2+), y16-98(2+), y22-98(3+), b3 |
| ELLIQFYK | | 527,302592 | 2 | y7, y6, y5, y4, y3, y6(2+), y5(2+), b3 |
| SGNIPAGTTVDTK | | 630,825149 | 2 | y10, y9, y8, y7, y6, y5, y4, y3, y10(2+), y9(2+), b3, b4 |
| SIEEQQKPLTDSQR | | 829,92084 | 2 | y10, y9, y8, y7, y5, y4, y3, y12, y11, b3, b4, b5, b6 |
| VELEVTLPGEGK | | 635,848092 | 2 | y11, y10, y9, y8, y7, y6, y5, y4, y3, b3, b4 |
| VLQPPSILYGGR | | 650,374612 | 2 | y11, y10, y9, y8, y7, y6, y5, y4, y3, y9(2+), y8(2+), b3 |

**Table S3. Quantification of phosphorylated peptides**

| Peptide Sequence and Modifications position | Phosphorylated residue | log2(LegK1/ eGFP) | LegK1/ eGFP | CI 2,5% | CI 97,5% | adjusted p-value |
|---|---|---|---|---|---|---|
| ARYHLVDKEHDS[Phospho]AEGSHTSGQSNGR | S824 | -0,594 | 0,66 | -0,89 | -0,29 | 7,15E-04 |
| SAS[Phospho]FNTDPYVR | S387 | 0,112 | 1,08 | -0,03 | 0,25 | 1,41E-01 |
| YHLVDKEHDS[Phospho] AEGS[Phospho]HTSGQSNGR | S824 & S828 | -0,756 | 0,59 | -1,40 | -0,11 | 5,18E-02 |
| YHLVDKEHDS[Phospho] AEGSHTSGQSNGR | S824 | -0,302 | 0,81 | -0,53 | -0,08 | 2,51E-02 |
| **Peptide Sequence and Modifications position** | **Phosphorylated residue** | **log2(LegK1-KA/eGFP)** | **LegK1-KA/eGFP** | **CI 2,5%** | **CI 97,5%** | **adjusted p-value** |
| ARYHLVDKEHDS[Phospho]AEGSHTSGQSNGR | S824 | -0,96 | 0,52 | -1,12 | -0,79 | 1,23E-17 |
| SAS[Phospho]FNTDPYVR | S387 | -0,19 | 0,88 | -0,42 | 0,03 | 1,37E-01 |
| YHLVDKEHDS[Phospho] AEGS[Phospho]HTSGQSNGR | S824 & S828 | -0,64 | 0,64 | -1,04 | -0,24 | 3,98E-03 |
| YHLVDKEHDS[Phospho] AEGSHTSGQSNGR | S824 | -0,61 | 0,65 | -0,88 | -0,35 | 3,52E-05 |
